# Genome-scale functional profiling of cell cycle controls in African trypanosomes

**DOI:** 10.1101/2020.07.16.206698

**Authors:** Catarina A. Marques, Michele Tinti, Andrew Cassidy, David Horn

## Abstract

Trypanosomatids, which include major pathogens of humans and livestock, are divergent eukaryotes for which cell cycle controls and the underlying mechanisms are not completely understood. Here, we describe a genome-wide RNA-interference library screen for cell cycle regulators in bloodstream form *Trypanosoma brucei*. We induced massive parallel knockdown and sorted the perturbed population into cell cycle stages using flow cytometry. RNAi-targets were deep-sequenced from each stage and cell cycle profiles were digitally reconstructed at a genomic scale. We identify hundreds of proteins that impact cell cycle progression; glycolytic enzymes required for G_1_S progression, DNA replication factors, mitosis regulators, proteasome and kinetochore complex components required for G_2_M progression, flagellar and cytoskeletal components required for cytokinesis, mRNA-binding factors, protein kinases and many previously uncharacterised proteins. The outputs facilitate functional annotation and drug-target prioritisation and provide comprehensive functional genomic evidence for the machineries, pathways and regulators that coordinate progression through the trypanosome cell cycle.

*The data can be searched and browsed using an interactive, open access, online data visualization tool (https://tryp-cycle.onrender.com).*

## Introduction

The canonical eukaryotic cell cycle encompasses discrete phases: G_1_ (gap 1), when the cell prepares for DNA replication; S (synthesis) phase, when nuclear DNA replication takes place; G_2_ (gap 2), when the cell prepares for mitosis; and M (mitosis) when the replicated DNA is segregated and the nucleus divides (1). Mitosis is immediately followed by cytokinesis (cell division), generating two daughter cells (2). Anomalies occurring during cell cycle progression can result in cell cycle arrest, to allow the cell to resolve the anomaly; in cell death, if the anomaly cannot be resolved or, among other outcomes, carcinogenesis.

Therefore, progression through the cell cycle is typically under strict checkpoint control; the G_1_-S, intra S phase, G_2_-M and spindle checkpoints control the onset of S phase, S phase progression, the onset of M phase and M phase progression, respectively (3). These processes have been extensively studied, particularly because cell cycle defects are common triggers for carcinogenesis (4). However, our understanding of the evolution and mechanisms of eukaryotic cell cycle progression control derives primarily from studies on the opisthokonts (including animals and fungi), with relatively fewer studies on more divergent eukaryotes, such as the trypanosomatids (1). The trypanosomatids are flagellated protozoa and include parasites that cause a range of neglected tropical diseases that have major impacts on human and animal health. The African trypanosome, *Trypanosoma brucei*, is transmitted by tsetse flies and causes both human and animal diseases, sleeping sickness and nagana, respectively, across sub-Saharan Africa (5). *T. brucei* has emerged as a highly tractable experimental system, both as a parasite and as a model organism (6). For example, the *T. brucei* flagellum (7) serves as a model for studies on human ciliopathies (8-11). Divergent features, shared with other pathogenic trypanosomatids, such as *Trypanosoma cruzi* and *Leishmania*, include glycolysis compartmentalised within glycosomes (12), a single mitochondrion with a complex mitochondrial DNA structure known as the kinetoplast (13) and polycistronic transcription of almost every gene (14). Widespread, and constitutive, polycistronic transcription in trypanosomatids places major emphasis on post-transcriptional controls by mRNA binding proteins (RBPs) and post-translational controls, involving protein phosphorylation in particular.

Studies that focus on cell cycle controls in *T. brucei* have revealed features that are conserved with other well-studied eukaryotes, but also features that are divergent (13,15). Most notably, the available evidence suggest that certain cell cycle checkpoints are absent. For example, cytokinesis is not dependent upon either mitosis or nuclear DNA synthesis in the insect stage of *T. brucei* (16). Moreover, functions previously thought to be fulfilled by highly conserved proteins employ lineage-specific or highly divergent proteins in trypanosomatids. The kinetochore complex, which directs chromosome segregation, is trypanosomatid-specific, for example (17), while the origin recognition complex (ORC), involved in DNA replication initiation, is highly divergent (18). In terms of high-throughput studies, transcriptome (19) and proteome (20) monitoring during the *T. brucei* cell cycle revealed hundreds of regulated mRNAs and proteins, while phosphoproteomic analysis revealed dynamic phosphorylation of several RBPs (21). Nevertheless, divergence presents a substantial challenge, many *T. brucei* genes have not yet been assigned a specific function, and many cell cycle regulators likely remain to be identified.

High-throughput functional genetic screens can be used to simultaneously assess every gene in a genome for a role in a particular process. We developed RNA interference Target Sequencing (RIT-seq) for *T. brucei* (22) and previously generated genome-scale fitness profiles, facilitating essentiality predictions and the prioritisation of potential drug targets (23). Here we describe a genome-scale RIT-seq screen to identify cell cycle controls and regulators in bloodstream form African trypanosomes. Following induction of knockdown, the cells were sorted according to their DNA content using fluorescence-activated cell sorting (FACS). The sorted populations were the G_1_, S and G_2_M cell cycle stages as well as perturbed cell populations with either less DNA than typically found in G_1_ or more DNA than typically found in G_2_M. RIT-seq analysis was then carried out for each sorted population and cell cycle profiles were digitally reconstructed for each gene using sequencing read-counts. This genome-wide screen reveals the protein complexes, pathways and signalling factors that coordinate progressive steps through the trypanosome cell cycle, both improving our understanding of trypanosome cell biology and also further facilitating the prioritisation of new potential drug targets.

## Results

### A genome-wide conditional knockdown screen for cell cycle progression defects

Bloodstream form *T. brucei* are readily grown in cell culture, with exponential proliferation and a doubling time of approximately 6.5 h. The *T. brucei* genome is diploid such that G_1_ cells have a 2C genome content; C represents the haploid DNA content. Cells progressing through nuclear S phase, and replicating their DNA, have a genome content between 2C and 4C, while cells that have completed DNA replication (G_2_M) have a 4C DNA content (***Figure 1A***). Cytokinesis then produces two daughter cells with a 2C DNA content. Some perturbations can yield defects which become apparent when cytokinesis produces sub-2C cells or over-replicated, polyploid (>4C) cells (***Figure 1A***). These polyploid cells arise due to endoreduplication, additional rounds of DNA replication without cytokinesis, either with (24) or without (25,26) mitosis, yielding cells with multiple diploid nuclei or cells with polyploid nuclei, respectively.

**Figure 1.**
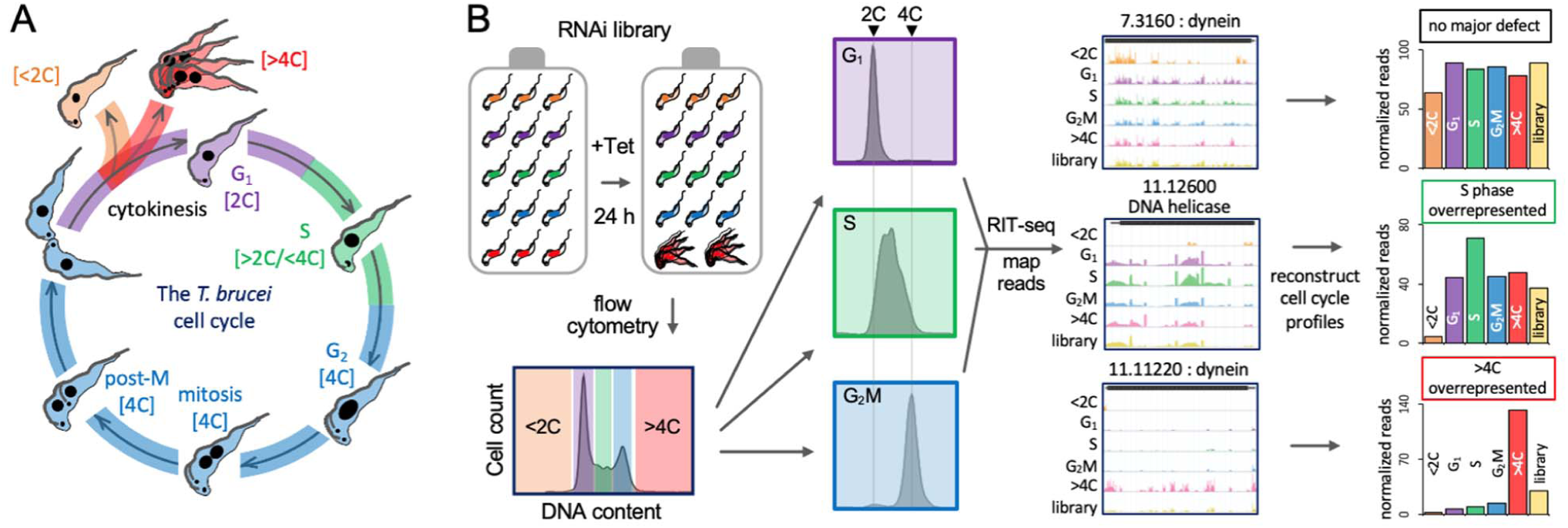
A genome-wide conditional knockdown screen for cell cycle progression defects. (**A**) Schematic representation of the bloodstream form *T. brucei* cell cycle, also showing aberrant sub-2C and >4C phenotypes. (**B**) The schematic illustrates the RIT-seq screen; massive parallel induction of RNAi followed by flow cytometry and RIT-seq, allowing for reconstruction of cell cycle profiles, using mapped reads from each knockdown. Each read-mapping profile encompasses the gene of interest and associated untranslated regions present in the cognate mRNA. The library data represents the uninduced and unsorted population. GeneIDs, Tb927.7.3160 for example, are shown without the common ‘Tb927.’ component.

We devised a high-throughput RNA interference (RNAi) Target sequencing (RIT-seq) screen to identify cell cycle controls and regulators at a genomic scale. Key features of RIT-seq screening include: first, use of a high-complexity *T. brucei* RNAi library comprising, in this case, approximately one million clones; second, massive parallel tetracycline-inducible expression of cognate dsRNA; and third, deep sequencing, mapping and counting of mapped reads derived from RNAi target fragments (22). Each clone in the library has one of approximately 100,000 different RNAi target fragments (250-1500 bp) between head-to-head inducible T7-phage promoters, with each cassette inserted at a chromosomal locus that supports robust expression. Inducibly expressed long dsRNA is then processed to siRNA by the native RNAi machinery (27). Complexity and depth of genome coverage in the library are critical, in that similar phenotypes produced by multiple RNAi target fragments against a single gene provide cross-validation. Improvements in reference genome annotation (28), next generation sequencing technology and sequence data analysis tools (see Materials and Methods) have also greatly facilitated quantitative phenotypic analysis using short-read sequence data.

Briefly, we induced massive parallel knockdown in an asynchronous *T. brucei* bloodstream form RNAi library for 24 h, fixed the cells, stained their DNA with propidium iodide (PI) and then used high-speed fluorescence-activated cell sorting (FACS) to divide the perturbed cell population into; sub-diploid (<2C), G_1_ (2C), S (between 2C and 4C), G_2_M (4C) and over-replicated (>4C) pools (***Figure 1—figure supplement 1***). Fixation and staining with the fluorescent DNA intercalating dye were pre-optimised for high-speed sorting (see Materials and Methods). Approximately 10 million cells were collected for each of the G_1_, S and G_2_M pools and samples from these pools were checked post-sorting to assess their purity (***Figure 1B, Figure 1—figure supplement 1***). For the perturbed and less abundant <2C and >4C pools, less than one million cells were collected; these pools were retained in their entirety for RIT-seq analysis.

RIT-seq was carried out for both the uninduced and induced, unsorted library controls, and for each of the five induced and sorted pools of cells as described in the Materials and Methods section. Briefly, we extracted genomic DNA from each sample, amplified DNA fragments containing each RNAi target fragment in PCR reactions (***Figure 1—figure supplement 1***) and used the amplified products to generate Illumina sequencing libraries. Analysis of sequencing reads mapped to the reference genome yielded counts for both total reads as well as reads containing the barcode (GTGAGGCCTCGCGA) that flanks each RNAi target fragment; the presence of the barcode confirmed that reads were derived from a specific RNAi target fragment and not from elsewhere in the genome. We derived counts of reads mapped to each of >7,200 non-redundant gene sequences in the uninduced and induced, unsorted library controls and in each of the five sorted samples. We selected the 24 h timepoint, equivalent to approximately 3.5 population doubling times, to allow sufficient time for the development of robust inducible phenotypes, but also to capture perturbed populations before they were critically diminished. Indeed, reads for 23.4% of genes dropped by >3-fold following 72 h of knockdown in a prior RIT-seq study (23), while a comparison of the unsorted control samples indicated that reads for only 0.6% of genes dropped by >3-fold following 24 h of knockdown here (see ***Figure 1—figure supplement 2***). Each sorted-sample library yielded between 22.6 and 37 million mapped read-pairs; <2C = 37 M, G_1_ = 35.1 M, S = 30.3 M, G_2_M = 22.6 M, >4C = 24.9 M; yielding >35,000 RNAi data-points (***Supplementary File 1***).

The RIT-seq digital data for individual genes following knockdown provided a measure of abundance in each pool and were, therefore, used to digitally reconstruct cell cycle profiles for individual gene knockdowns (***Figure 1B***). We expected to observe accumulation of particular knockdowns in specific cell cycle phase pools, thereby reflecting specific defects. This was indeed the case, and some examples are shown to illustrate; no major defect, S phase overrepresented or >4C overrepresented, following knockdown (***Figure 1B***). These outputs suggest that loss of the cytoplasmic dynein heavy chain (Tb927.7.3160) does not perturb cell cycle distribution; that a putative DNA helicase (Tb927.11.12600) is required for the completion of S phase; and that knockdown of the axonemal dynein heavy chain (Tb927.11.11220) results in endoreduplication in the absence of cytokinesis. Dyneins are cytoskeletal motor proteins that move along microtubules, to produce a flagellar beat, for example (29).

### Validation and identification of >1,000 candidates linked to cell cycle defects

The *T. brucei* core genome comprises a non-redundant set of over 7,200 protein-coding sequences, for which we were now able to digitally reconstruct cell cycle profiles following knockdown; the full dataset is shown mapped to the eleven megabase chromosomes in ***Figure 2—figure supplement 1***. The data can also be searched and browsed using an interactive, open access, online data visualization tool (see ***Figure 2— figure supplement 2***; https://tryp-cycle.onrender.com). First, we examined knockdowns reporting an overrepresentation of >4C cells, indicating endoreduplication, and this yielded 201 genes for which reads in the >4C pool exceeded 1.5-fold the sum of reads in the G_1_, S phase and G_2_M pools (***Figure 2A***, left-hand panel; ***Supplementary File 1***). The >4C phenotype was previously observed following α-tubulin knockdown in a landmark study that first described RNAi in *T. brucei* (24) and, indeed, we observed pronounced overrepresentation of >4C cells for both adjacent α-tubulin (Tb927.1.2340) and β-tubulin (Tb927.1.2350) gene knockdowns (***Figure 2A***, middle and right-hand panel). We then examined knockdowns reporting an overrepresentation of <2C cells, indicating a reduced DNA content. This yielded 119 genes for which reads in the <2C pool exceeded 1.5-fold the sum of reads in the G_1_, S phase and G_2_M pools (***Figure 2B***, left-hand panel; ***Supplementary File 1***). Haploid cells were previously observed following DOT1A knockdown (30) and, consistent with the previous report, we observed pronounced overrepresentation of <2C cells for the DOT1A (Tb927.8.1920) gene knockdown (***Figure 2A***, middle and right-hand panel); we are not aware of other knockdowns reported to yield a similar phenotype. Together, these results provide initial validation for the >4C and <2C components of the screen.

**Figure 2.**
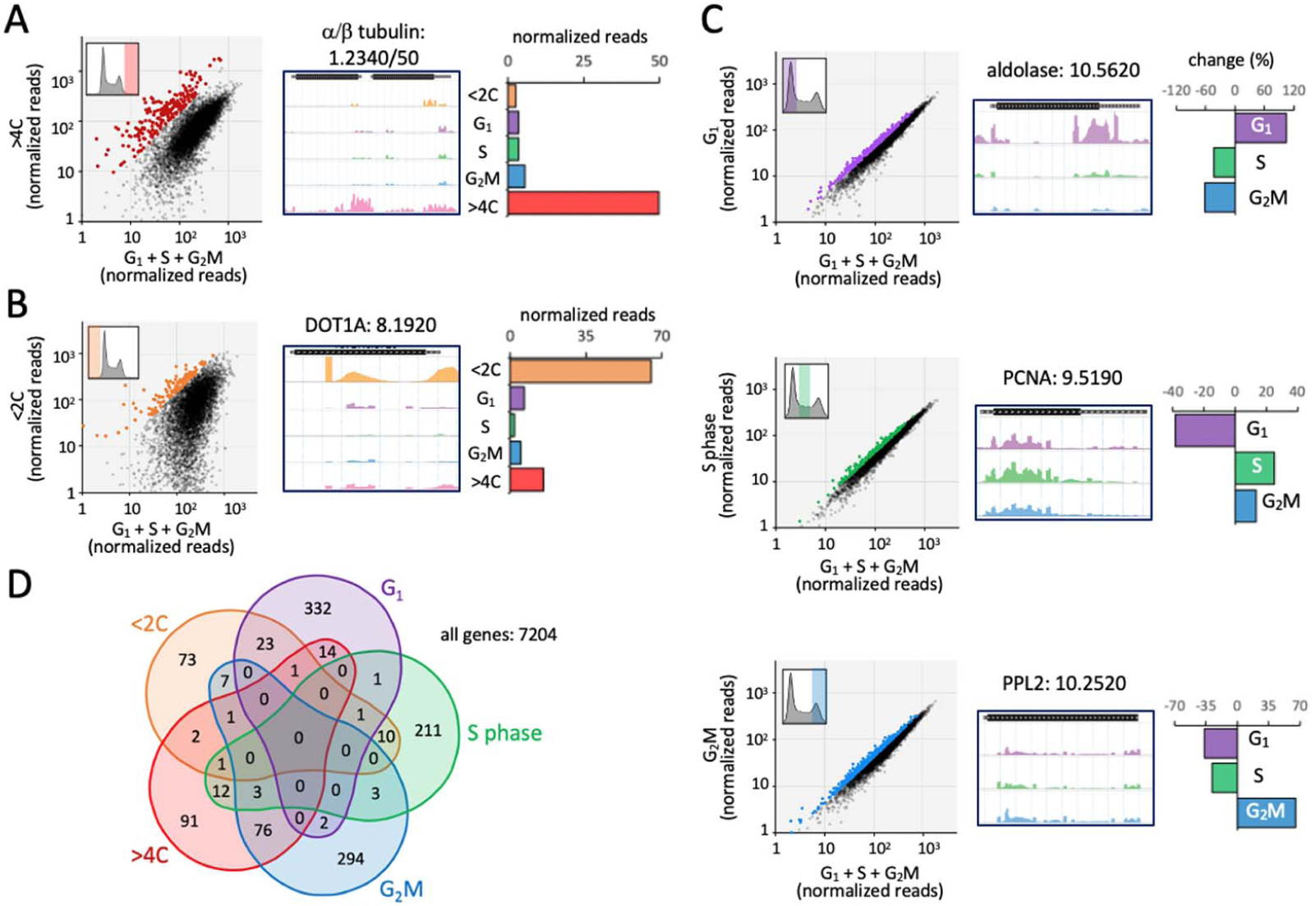
Validation and identification of >1000 candidates linked to cell cycle defects. (**A**) The plot on the left shows knockdowns overrepresented in the >4C experiment in red; those with >1.5-fold the sum of reads in the G_1_, S phase and G_2_M samples combined. The read-mapping profile and read-counts for α/β-tubulin are shown to the right. (**B**) The plot on the left shows knockdowns overrepresented in the sub-2C experiment in orange; those with >1.5-fold the sum of reads in the G_1_, S phase and G_2_M samples combined. The read-mapping profile and read-counts for DOT1A are shown to the right. (**C**) The plots on the left shows knockdowns overrepresented in the G_1_, S phase and G_2_M experiments in purple, green and blue, respectively; those that were >25% overrepresented in each category. Read-mapping profiles and relative read-counts for example hits are shown to the right. PCNA, proliferating cell nuclear antigen; PPL2, PrimPol-like 2. (**D**) The Venn diagram shows the distribution of knockdowns overrepresented in each arm of the screen.

Next, we turned our attention to knockdowns reporting an overrepresentation of G_1_, S phase or G_2_M cells. The pools of knockdowns that registered >25% overrepresented read counts in each of these categories are highlighted in ***Figure 2C*** (left-hand panels and ***Supplementary File 1***) and data for an example from each category are shown (***Figure 2C***; right-hand panels); the glycolytic enzyme, aldolase (Tb927.10.5620), reported 104% increase in G_1_ cells (further details below); the proliferating cell nuclear antigen (PCNA; Tb927.9.5190), a DNA sliding clamp that is a central component of the replication machinery (31), reported 25% increase in S phase cells, consistent with prior analysis (32); and PrimPol-like 2 (PPL2; Tb927.10.2520), a post-replication translesion polymerase, reported 65% increase in G_2_M cells, also consistent with prior analysis (33). These results provided initial validation for the G_1_, S phase and G_2_M components of the screen.

Overall, the five components of the screen yielded 1,158 genes that registered a cell cycle defect, based on the thresholds applied above. This is 16.1% of the 7,204 genes analysed, and the distribution of these genes among the five arms of the screen are shown in the Venn diagram in ***Figure 2D***. Since we predicted that knockdowns associated with a cell cycle defect were more likely to also register a growth defect, we compared these datasets to prior RIT-seq fitness profiling data (23). All groups of genes that registered cell cycle defects, except for the <2C set, were significantly enriched for genes that previously registered a loss-of-fitness phenotype following knockdown in bloodstream form cells (χ^2^ test; <2C, *p* = 0.15; G_1_, *p* = 0.015; S, *p* = 4.7^-4^; G_2_M, *p* = 3.5^-24^; >4C, *p* = 4.4^-199^). This is consistent with loss-of-fitness as a common outcome following a cell cycle progression defect. Taken together, the analyses above provided validation for the RIT-seq based cell cycle phenotyping approach and yielded >1,000 candidate genes that impact specific steps during *T. brucei* cell cycle progression.

### Cytokinesis defects associated with endoreduplication

In bloodstream form *T. brucei*, defective >4C cells can arise due to endoreduplication without cytokinesis, either with (24) or without (25,26) mitosis. Cytokinesis-only defects were previously observed following knockdown of α-tubulin (24) or flagellar proteins (7,34); flagellar beat is thought to be required for cytokinesis in bloodstream form *T. brucei*.

Consistent with these observations, α-tubulin (see ***Figure 2A***) and axonemal dynein heavy chain (see ***Figure 1B***) knockdowns were amongst 201 genes overrepresented in the >4C pool in our screen. Gene Ontology (GO) annotations provide structured descriptions of gene products in terms of functions, processes, and compartments. Analysis of overrepresented GO annotations within the >4C-enriched cohort revealed ‘dynein’, ‘intraflagellar transport’ (IFT), ‘axoneme’ and ‘cytoskeleton’ terms, and also ‘chaperonin T-complex’, ‘cytokinesis’ and multiple other ‘cell cycle’ terms (***Figure 3A***). The violin plot in ***Figure 3B*** shows specific enrichment of IFT and dynein knockdowns in the >4C pool, relative to other cohorts of knockdowns. Exocyst components, primarily involved in exocytosis (35), were included as a negative control cohort; indeed, none of the exocyst components register enrichment in the >4C pool, nor in any other pool (see below). Enrichment of individual chaperonin T-complex components, dyneins, and IFT factors in the >4C pool is illustrated in ***Figure 3C***. The chaperonin T-complex is involved in tubulin and actin folding (36) and, notably, actin knockdown was also associated with >4C enrichment (***Figure 3—figure supplement 1***).

**Figure 3.**
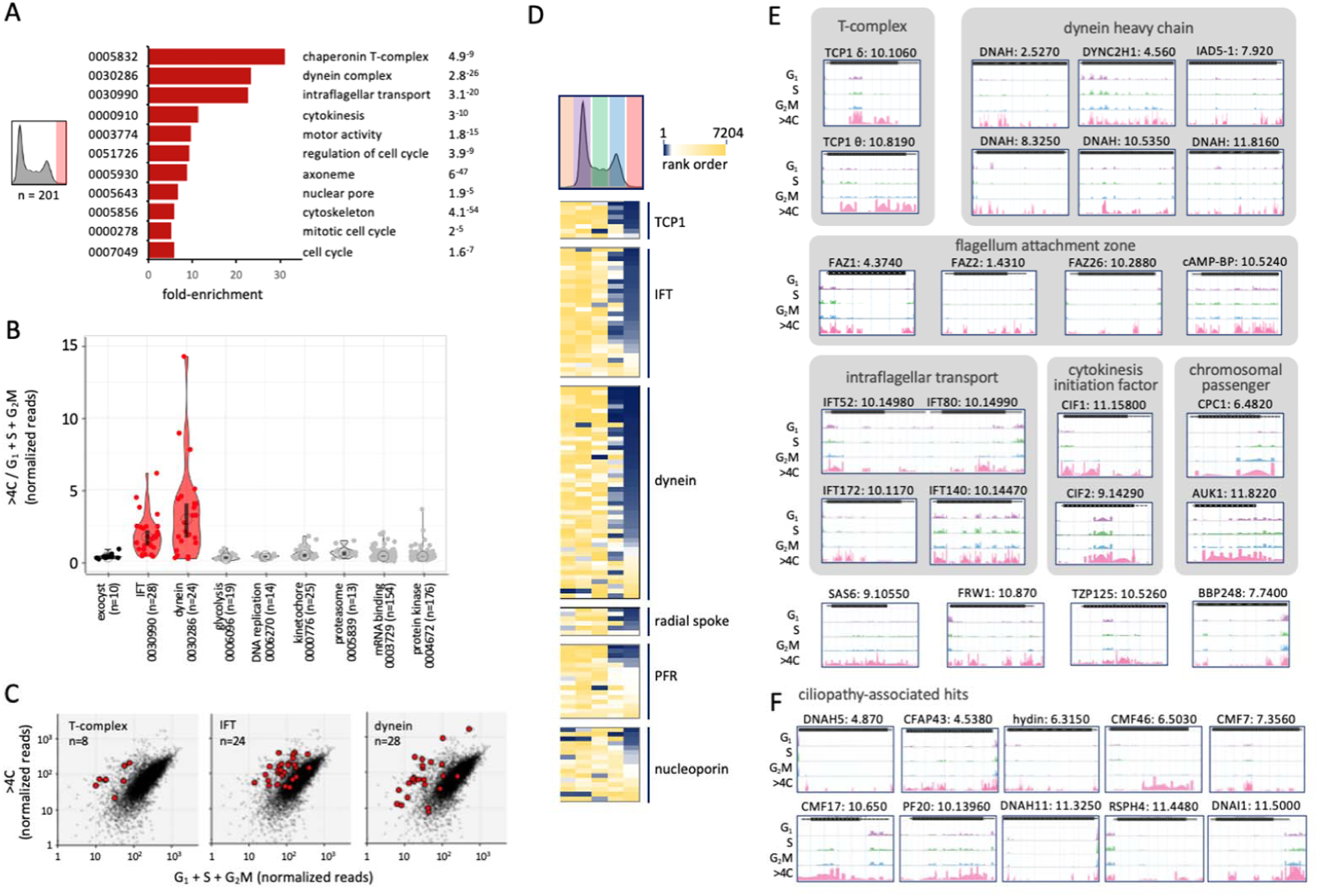
Cytokinesis defects associated with endoreduplication. (**A**) The bar-graph shows enriched Gene Ontology terms in the >4C overrepresented dataset. (**B**) The violin plot shows relative >4C read-counts for cohorts of genes and reflects data distribution. Open circles indicate median values and the vertical bars indicate 95% confidence intervals. Significantly overrepresented cohorts are indicated in red. (**C**) The plots show overrepresentation of T-complex, dynein and intraflagellar transport (IFT) factors in red in the >4C experiment. (**D**) The heatmaps show relative representation in all five sorted pools for the above and additional cohorts of knockdowns; blue, most overrepresented. (**E**) Example read-mapping profiles for hits overrepresented in the >4C pool. (**F**) Example read-mapping profiles for ciliopathy-associated hits overrepresented in the >4C pool. CMF, Component of Motile Flagella; CFAP, Cilia and Flagella Associated Protein.

The heat-map in ***Figure 3D*** shows the data for all five sorted pools for the cohorts described above and for additional cohorts of knockdowns enriched in the >4C pool; these include radial spoke proteins, extra-axonemal paraflagellar rod (PFR) proteins, as well as nucleoporins. The gallery in ***Figure 3E*** shows examples of RIT-seq read-mapping profiles for twenty-four individual genes that register >4C enrichment following knockdown. In addition to the categories above, these include the inner arm dynein 5-1 (37), FAZ proteins which mediate attachment of the flagellum to the cell body (38); cytokinesis initiation factors CIF1/TOEFAZ1 (Tb927.11.15800) and CIF2 (Tb927.9.14290) (39) and chromosomal passenger complex components, including CPC1 (Tb927.6.4820) and the aurora B kinase, AUK1 (Tb927.11.8220). AUK1 and CPC1 are spindle-associated and regulate mitosis and cytokinesis (26,40). Notably, endoreduplication was reported previously following AUK1 knockdown in bloodstream form *T. brucei* (41) and this is the kinase with the most pronounced overrepresentation in our >4C dataset. The next >4C overrepresented kinase is the CMGC/RCK (Tb927.3.690), knockdown of which previously yielded a dramatic cytokinesis defect (42).

Additional examples of genes registering >4C overrepresentation include the centriole cartwheel protein SAS6 (Tb927.9.10550) (43), the cleavage furrow-localizing protein FRW1 (Tb927.10.870) (44), the basal body – axoneme transition zone protein TZP125 (Tb927.10.5260) (45) and the basal body protein BBP248 (Tb927.7.7400) (46). One hundred additional examples are shown in ***Figure 3—figure supplement 1***, including intermediate and light chain dyneins; other flagellum-associated factors, radial spoke proteins, components of motile flagella, flagellum attachment and transition zone proteins, intraflagellar transport proteins, kinesins (47,48), nucleoporins (49), and many previously uncharacterised hypothetical proteins. Some other notable examples include the microtubule-severing katanin KAT80 (Tb927.9.9960) (50), the dynein regulatory factor trypanin (Tb927.10.6350) (51), the AIR9 microtubule associated protein (Tb927.11.17000) (52), CAP51V (Tb927.7.2650) (53) and importin, IMP1 (Tb927.9.13520) (54).

Orthologues of several *T. brucei* flagellar proteins have previously been linked to debilitating human ciliopathies, such that the trypanosome flagellum is used as a model for studies on these defects. Defects in intraflagellar dynein transport are associated with respiratory infections, for example (9). Some examples of ciliopathy-associated orthologues which register overrepresentation in the >4C pool are shown in ***Figure 3F*** and ***Figure 3— figure supplement 1***. These include proteins linked to primary ciliary dyskinesia (DNAI1, Tb927.11.5000; DNAH5, Tb927.4.870; DNAH11, Tb927.11.3250; RSPH4, Tb927.11.4480) (11); male infertility (CFAP43, Tb927.4.5380; CMF7/TbCFAP44, Tb927.7.3560) (10); and cone-rod dystrophies, as well as other ocular defects (CMF17, Tb927.10.650; CMF39, Tb927.4.5370; CMF46, Tb927.6.5030) (8).

From analysis of knockdowns overrepresented in the >4C pool, we conclude that RIT-seq screening provided comprehensive genome-scale identification of cytokinesis defects in bloodstream form *T. brucei*. Endoreduplication appears to be a common outcome following a cytokinesis defect. Amongst hundreds of genes required for progression through cytokinesis, flagellar proteins featured prominently, including the majority of dynein chains and intraflagellar transport factors. Many of these factors are essential for viability and include potential druggable targets in trypanosomatids, as well as orthologues of proteins associated with ciliopathies.

### Defects producing sub-diploid cells

A DNA replication or mitosis defect followed by cytokinesis may result in generation of cells that retain nuclear DNA with a sub-2C DNA content. We emphasise retention of nuclear DNA here because *T. brucei* cells lacking nuclear DNA, referred to as zoids, have been reported previously as a result of asymmetrical cell division. Zoids are typically observed when DNA replication or mitosis are perturbed in insect stage cells (16,25,55). The zoid phenotype is typically either absent or less abundant in the developmentally distinct bloodstream form cells (56) that we analysed here. Nevertheless, any zoids present in the <2C pool will not have been detected using RIT-seq, since detection relies upon the presence of a nuclear RNAi target fragment.

One hundred and nineteen knockdowns were overrepresented in the <2C RIT-seq screening dataset. A particularly prominent hit was the previously identified histone methyltransferase, DOT1A (see above). DOT1A is responsible for dimethylation of histone H3K76, and DOT1A knockdown results in premature mitosis without DNA replication, generating cells with a haploid DNA content (30). The gallery in ***Figure 4*** shows examples of RIT-seq read-mapping profiles for twelve additional genes that register <2C enrichment following knockdown. These include the DNA replication origin recognition complex-associated protein, ORC1B (Tb927.9.2030) (57), two putative protein kinases (NEK7, Tb927.3.3190; CDPK, Tb927.4.3770), a putative mRNA-binding helicase (ZC3H41, Tb927.11.1980), a nucleosome assembly protein (NAP, Tb927.3.4880) and threonyl-tRNA synthetase (Tb927.5.1090). Notably, threonine metabolism impacts histone methylation in mammalian cells (58), while a protein containing a putative methyllysine-binding SET domain (Tb927.6.910) also registers enrichment in the <2C pool. These hits present new candidate regulators of DNA replication, mitosis or meiosis (59), and further potential links to post-translational protein methylation as a key player in coordinating these processes.

**Figure 4.**
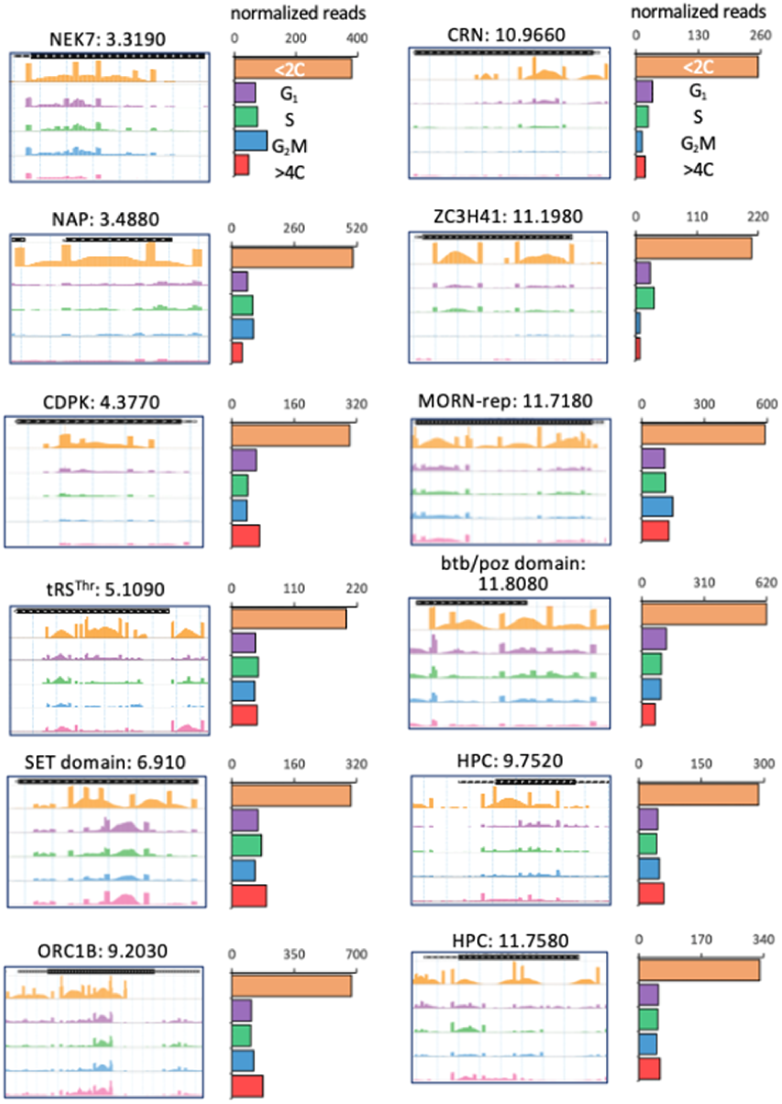
Defects producing sub-diploid cells. Read-mapping profiles and read-counts for example hits overrepresented in the <2C experiment. HPC, Hypothetical Protein, Conserved.

### A profile of G_1_, S phase and G_2_M defects

We next analysed knockdowns overrepresented in the G_1_, S phase or G_2_M pools. Several hundred knockdowns registered >25% overrepresented read counts in each of these categories, as shown in a RadViz plot (***Figure 5A***). GO annotations within each cohort revealed a number of enriched terms (***Figure 5B***). Overrepresented knockdowns were associated with glycolysis and mRNA binding in the G_1_ pool, with DNA replication in the S phase pool and with a similar profile to that seen for the >4C set in the G_2_M pool.

**Figure 5.**
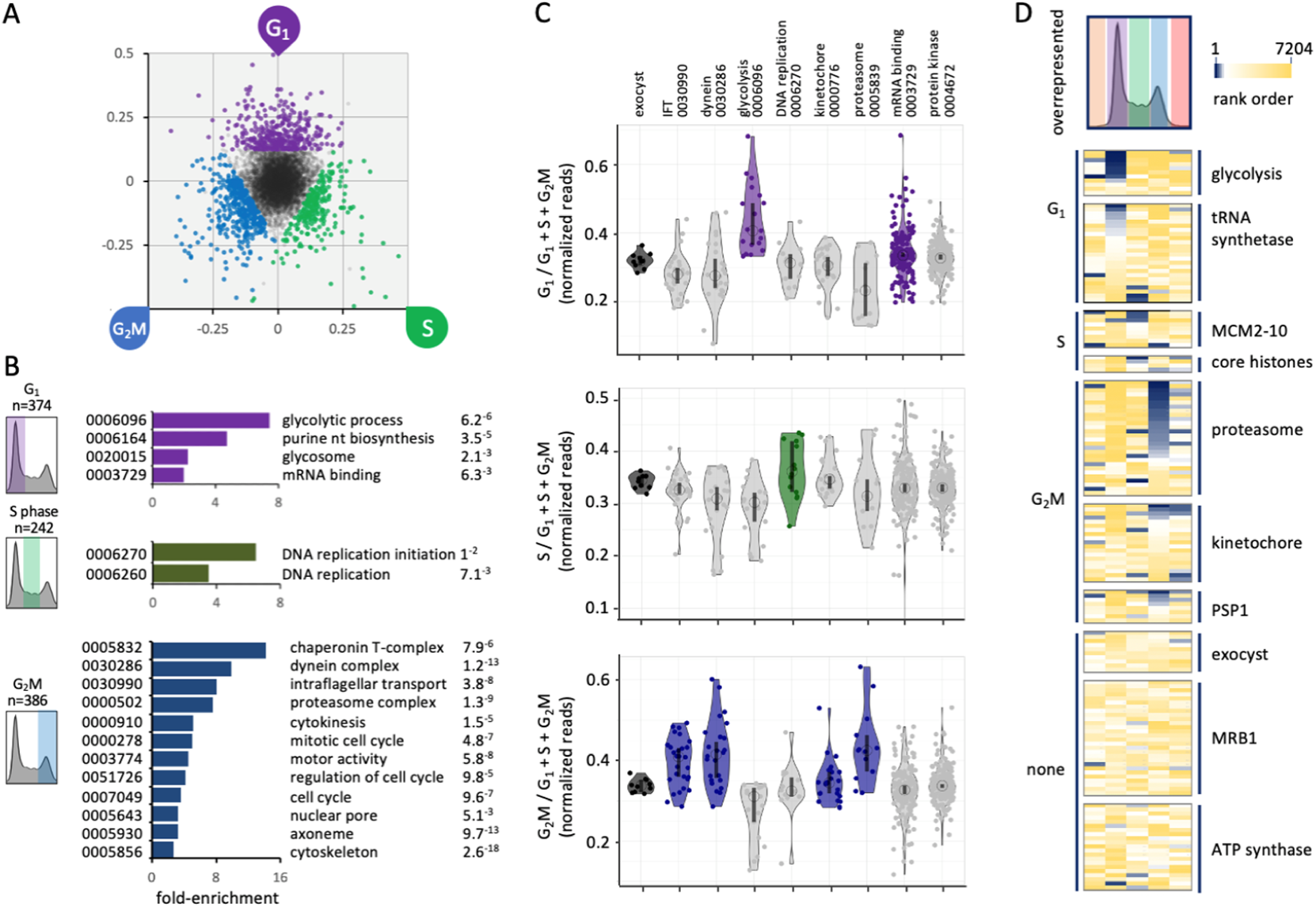
A profile of G_1_, S phase and G_2_/mitosis defects. (**A**) The RadViz plot shows knockdowns that registered >25% overrepresented read-counts in the G_1_, S phase or G_2_M categories. (**B**) The bar-graphs shows enriched Gene Ontology terms in the G_1_, S phase or G_2_M overrepresented datasets. (**C**) The violin plots show relative G_1_, S phase or G_2_M read-counts for cohorts of genes and reflects data distribution. Open circles indicate median values and the vertical bars indicate 95% confidence intervals. Overrepresented cohorts are indicated in purple, green and blue, respectively. (**D**) The heatmaps show relative representation in all five sorted pools for the above and additional cohorts of knockdowns; blue, most overrepresented.

The violin plot in ***Figure 5C*** shows specific enrichment of individual knockdowns for glycolytic enzymes and a subset of mRNA binding protein in the G_1_ pool, for DNA replication factors in the S phase pool, and proteasome components and a subset of kinetochore component in the G_2_M pool (***Figure 5C***, lower panel). Overlap between knockdowns that accumulate in both the G_2_M and >4C pools likely reflects mitosis and cytokinesis defects both before and after endoreduplication; perhaps 24 h of knockdown is insufficient for endoreduplication in all perturbed cells or not all mitosis or cytokinesis-perturbed phenotypes result in endoreduplication; compare >4C and G_2_M data for IFT factors and dyneins in ***Figure 3B*** and ***Figure 5C***, for example. Once again, exocyst components provided a control cohort with no components registering enrichment in the G_1_, S phase or G_2_M pools following knockdown (***Figure 5C***).

The heat-map in ***Figure 5D*** shows the data for all five sorted pools for the cohorts described above and for additional knockdowns enriched in the G_1_ or S phase (tRNA synthetases), S phase (core histones) or G_2_M pools (PSP1, DNA polymerase suppressor 1), or not enriched in any pool. These latter sets provide further controls that do not appear to have specific impacts on cell cycle progression, including the mitochondria RNA editing accessory complex MRB1 (60) and the mitochondrial ATP synthase complex V (61). Thus, we identify a number of protein complexes, pathways and regulatory factors that are specifically required for progressive steps through the trypanosome cell cycle.

### Pathways and protein complexes associated with G_1_, S phase and G_2_M defects

We next explored some of the cohorts of hits described above in more detail. Glycolytic enzymes are particularly prominent amongst knockdowns that accumulate in G_1_ and we illustrate the RIT-seq profiling data for these enzymes in ***Figure 6A***. Seven of eleven glycolytic enzyme knockdowns register >25% overrepresentation in the G_1_ pool; hexokinase (Tb927.10.2010), phosphofructokinase (Tb927.3.3270), aldolase (see ***Figure 2C***), triosephosphate isomerase (Tb927.11.5520), glyceraldehyde 3-phosphate dehydrogenase (Tb927.6.4280), phosphoglycerate kinase C (Tb927.1.700) and pyruvate kinase (Tb927.10.14140). Glycolysis operates in peroxisome-like organelles known as glycosomes in trypanosomes and is thought to be the single source of ATP in bloodstream form cells (12). Glycolysis also provides metabolic intermediates that support nucleotide production. Notably, mammalian cell proliferation is accompanied by activation of glycolysis, and the Warburg effect relates to this phenomenon in oncology (62). Indeed, hexokinase regulates the G_1_/S checkpoint in tumour cells (63). The results are also consistent with the observation that *T. brucei* accumulate in G_1_ or G_0_ under growth-limiting conditions (64) or during differentiation to the non-dividing stumpy form (65), possibly reflecting a role for glucose sensing in differentiation (66). Notably, glycolytic enzymes are downregulated 6.7+/-5.2-fold in stumpy-form cells (67). We conclude that, as in other organisms (68), there is metabolic control of the cell cycle and a nutrient sensitive restriction point in *T. brucei*, with glycolysis playing a role in the G_1_ to S phase transition.

**Figure 6.**
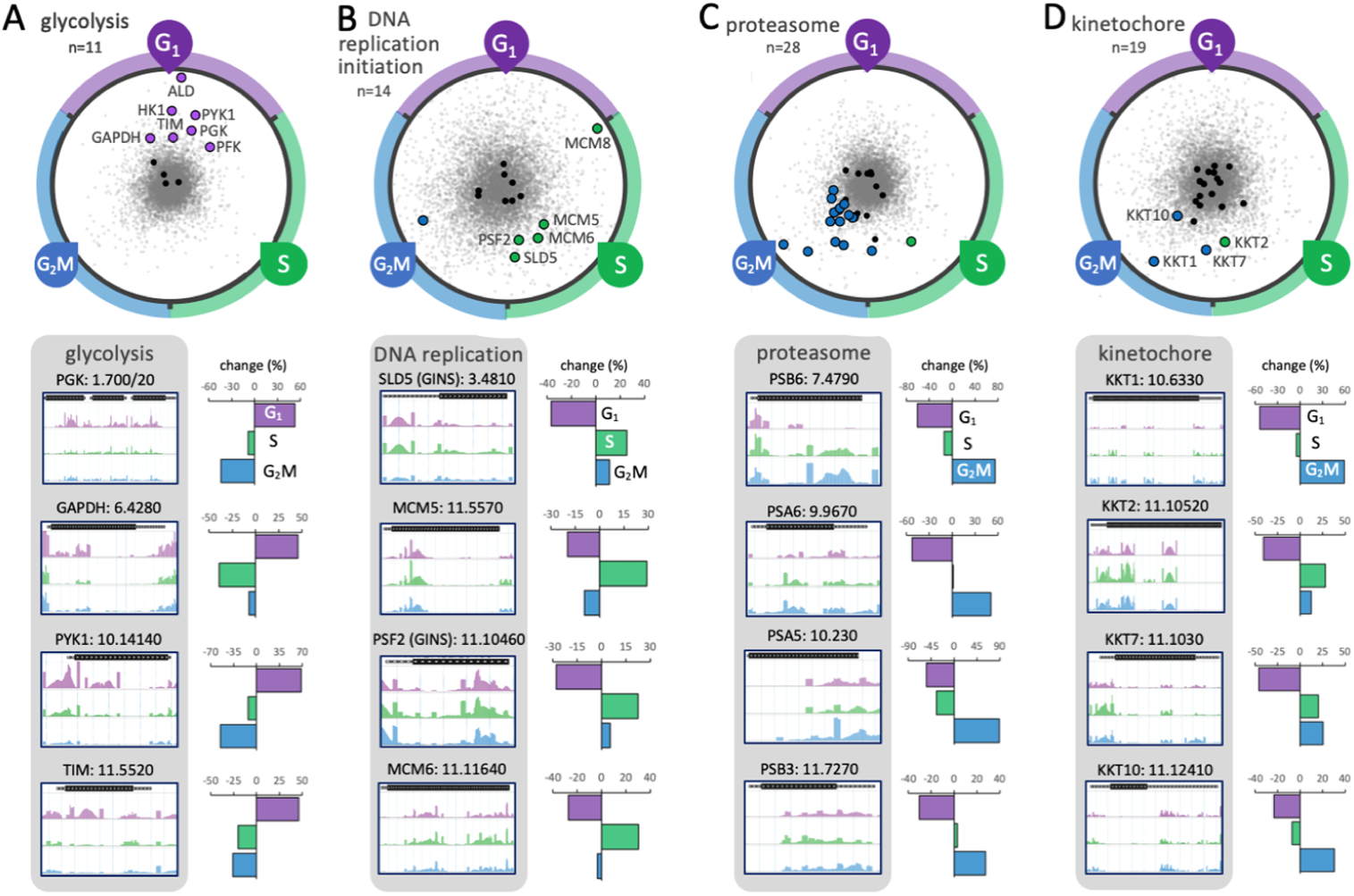
Protein complexes and pathways associated with G_1_, S phase and G_2_/mitosis defects. (**A**) The RadViz plot shows glycolytic enzyme knockdowns. Those that registered >25% overrepresented read-counts in the G_1_ category are indicated, purple. Black data-points indicate other genes from each cohort. Grey data-points indicate all other genes. The read-mapping profiles and relative read-counts in the lower panel show example hits. (**B**) As in a but for DNA replication initiation factor knockdowns that registered >25% overrepresented read-counts in the S phase category, indicated in green. (**C**) As in a but for proteasome component knockdowns that registered >25% overrepresented read-counts in the G_2_M category, indicated in blue. (**D**) As in a but for kinetochore component knockdowns that registered >25% overrepresented read-counts in the S phase or G_2_M categories, indicated in green or blue, respectively.

DNA replication initiation factors are particularly prominent amongst knockdowns that accumulate in S phase and we illustrate the RIT-seq profiling data for these factors in ***Figure 6B***. Five knockdowns that register >25% overrepresentation in the S phase pool are components of the eukaryotic replicative helicase, the CMG (Cdc45-MCM-GINS) complex.

At the core of this complex is the minichromosome maintenance complex (MCM2-7), a helicase that unwinds the duplex DNA ahead of the moving replication fork (69). Identification of this subset of components suggests that these particular subunits are limiting for progression through S phase. Proteasome components are particularly prominent amongst knockdowns that accumulate in G_2_M, and we illustrate the RIT-seq profiling data for this protein complex in ***Figure 6C***. Sixteen of 28 proteasome component knockdowns register >25% overrepresentation in the G_2_M pool. This output suggests that the *T. brucei* proteasome is most likely responsible for degrading cell cycle regulators, such as poly-ubiquitinated cyclins, which are known to control cell cycle checkpoints in other eukaryotes. Candidate target cyclins in *T. brucei* include: cyclin 6 (CYC6, Tb927.11.16720), degradation of which is required for mitosis (70); cyclin-like CFB2 (Tb927.1.4650), required for cytokinesis (71); cyclin 2 (CYC2, Tb927.6.1460) or cyclin 3 (CYC3, Tb927.6.1460), which have short half-lives and a candidate destruction box motif in the case of CYC3 (72).

Kinetochore components (17) are also amongst knockdowns that accumulate in G_2_M and we illustrate the RIT-seq profiling data for this protein complex in ***Figure 6D***. Although knockdown of KKT2 (Tb927.10.10520), a putative kinase, registered overrepresentation in the S phase pool, KKT1 (Tb927.10.6330), KKT7 (Tb927.11.1030) and KKT10 (CLK1, Tb927.11.12410) knockdowns registered >25% overrepresentation in the G_2_M pool, suggesting that these particular kinetochore components, which all display temporal patterns of phosphorylation from S phase to G_2_M (21), are limiting for progression through mitosis. Notably, KKT10 is a kinase responsible for phosphorylation of KKT7, which is required for the metaphase to anaphase transition (73); as well as for the phosphorylation of KKT1, which is required for kinetochore assembly (74). These findings are consistent with the view that kinetochore components control a non-canonical spindle checkpoint in trypanosomes (73).

### RBPs, kinases and hypothetical proteins associated with G_1_, S phase and G_2_M defects

Widespread polycistronic transcription in trypanosomatids places great emphasis on post-transcriptional controls and, consistent with this, knockdowns overrepresented in the G_1_, S phase and G_2_M pools revealed many putative mRNA binding proteins (RBPs) and kinases. Indeed, RBPs are significantly enriched amongst knockdowns that registered G_1_, S phase or G_2_M cell cycle defects (χ^2^ test, *p* = 7^-5^). We show the RIT-seq profiling data for seven RBP knockdowns that register >25% overrepresentation in these pools (***Figure 7A***). These include knockdowns for two components of the translation initiation factor, eIF3, linked to accumulation in G_1_ (Tb927.8.1170; Tb927.10.8270). Proposed regulatory functions for eIF3 include reinitiation of translation on polycistronic mRNAs and as a substrate for translation regulatory kinases (75). The RBP10 (Tb927.8.2780), RBP29 (Tb927.10.13720) and ZC3H31 (Tb927.10.5150) knockdowns were also enriched in G_1_. RBP10, in particular, has been characterised in some detail and promotes the bloodstream form state (76). Thus, the RIT-seq cell cycle screen implicated a number of specific RBPs in post-transcriptional control of cell cycle progression, most likely through modulation of mRNA stability and/or translation of cell cycle regulators.

**Figure 7.**
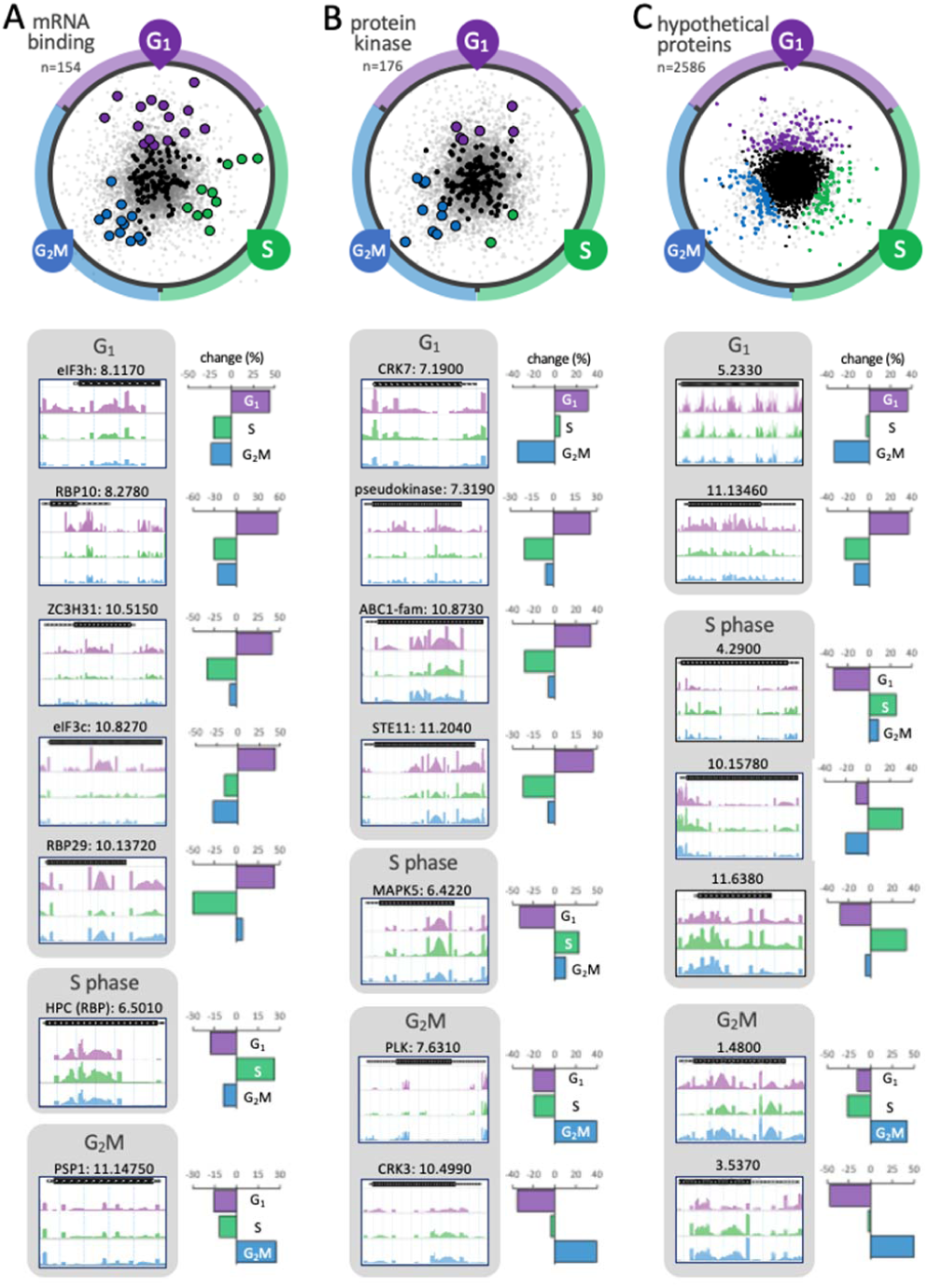
RBPs kinases and hypothetical proteins associated with gap and S phase defects. (**A**) The RadViz plot shows mRNA binding protein knockdowns (RBPs). Those that registered >25% overrepresented read-counts in the G_1_, S phase or G_2_M categories are indicated, in purple, green and blue, respectively. The read-mapping profiles and relative read-counts in the lower panels show example hits. (**B**) As in a but for protein kinase knockdowns. (**C**) As in a but for hypothetical (conserved) protein knockdowns.

We show data for several protein kinases above, linked to enriched >4C (***Figure 3D***), sub-2C (***Figure 4***), S phase or G_2_M (***Figure 6D***) phenotypes, and now show the RIT-seq profiling data for seven additional protein kinase knockdowns that register >25% overrepresentation in the G_1_, S phase or G_2_M pools (***Figure 7B***). These include knockdowns for CRK7 (Tb927.7.1900), linked to accumulation in G_1_; MAPK5 (Tb927.6.4220), linked to accumulation in S phase and polo-like kinase (PLK, Tb927.7.6310) and cdc2-related kinase 3 (CRK3, Tb927.10.4990), linked to accumulation in G_2_M. PLK was previously shown to control cell morphology, furrow ingression and cytokinesis (77-79), while CRK3 was shown to play a role in G_2_M progression in bloodstream form *T. brucei* (42,80).

Finally, we analysed genes encoding proteins annotated as hypothetical (conserved). Despite excellent progress in genome annotation, 35% of the non-redundant gene-set in *T. brucei* retain this annotation, amounting to >2,500 genes. We show data for more than twenty of these knockdowns above, linked to enriched >4C (***Figure 3—figure supplement 1***) or sub-2C (***Figure 4***) phenotypes, and we identify additional hypothetical (conserved) protein knockdowns that register >25% overrepresentation in the G_1_, S phase or G_2_M pools RIT-seq profiling data are shown for seven examples in ***Figure 7C*** and for twenty four examples in ***Figure 7 – figure supplement 1***. Amongst fifty-four other examples of knockdowns shown in ***Figure 7—figure supplement 1***, are alternative oxidase (Tb927.10.7090) (81), linked to G_1_ enrichment; tRNA synthetases linked to G_1_ (Trp, Tb927.3.5580; His, Tb927.6.2060) or S phase enrichment (Glu, Tb927.6.4590; Lys, Tb927.8.1600); kinesins linked to S phase (Tb927.7.7120) or G_2_M enrichment, including both chromosomal passenger complex kinesins (26) (Tb927.11.2880, KIN-A and Tb927.7.5040, KIN-B) and KIN-G (Tb927.6.1770); CYC6 (25,82), CFB2 (71), centrin 3 (Tb927.10.8710) (83) and, finally, both components of the histone chaperone FACT (facilitates chromatin transcription) complex (84) Spt16 (Tb927.3.5620) and Pob3 (Tb927.10.14390), linked to G_2_M enrichment. Notably, the FACT complex has been linked to centromere function in human cells (85).

### Linking cell cycle regulated transcripts and proteins to cell cycle progression defects

Some factors required for cell cycle progression are themselves cell cycle regulated.

To identify some of these factors, we compared our functional assignment data with quantitative transcriptome (19), proteome (20) and phosphoproteome (21) cell cycle profiling data. An initial survey of all 1,158 genes that registered a cell cycle defect here (see ***Figure 2D***) revealed significant enrichment of cell cycle regulated mRNAs (overlap = 105 of 485, χ^2^ *p* = 8.5^-4^), as well as proteins displaying cell cycle regulated phosphorylation (overlap = 106 of 547, χ^2^ *p* = 0.035). It is important to note here, however, and for the analyses below, that these transcriptome and (phospho)proteome datasets were derived from insect stage *T. brucei*, meaning that regulation may differ in some cases in bloodstream *T. brucei* cells used for RIT-seq analysis here.

Specific transcripts required for cell cycle progression are likely upregulated prior to peak demand for the encoded protein, and we found evidence to support this view. For example, transcripts upregulated in late G_1_ or in S phase were enriched amongst those knockdowns linked to accumulation in the G_2_M pool here (χ^2^ *p* = 3.3^-3^ and *p* = 1.1^-2^ respectively); G_1_ upregulated transcripts included, for example, both components of the FACT complex (see ***Figure 7—figure supplement 1***). In addition, S phase and G_2_M upregulated transcripts, including those encoding multiple flagellum-associated proteins, were enriched amongst knockdowns linked to accumulation in the >4C pool (χ^2^ *p* = 4.6^-18^ and *p* = 2.4^-5^ respectively).

Although the overlap between cell cycle regulated proteins (20) and those knockdowns that registered a cell cycle defect here failed to achieve significance (overlap = 71 of 367, χ^2^ *p* = 0.09), cell cycle regulated proteins do appear to be required for progression through specific stages of the cell cycle. For example, multiple glycolytic enzymes upregulated in G_1_ were linked to accumulation in the G_1_ pool following knockdown (χ^2^ *p* = 7.9^-11^). In addition, proteins highly upregulated in G_2_ and M were linked to accumulation in the G_2_M (χ^2^ *p* = 1.8^-8^) or >4C pools (χ^2^ *p* = 8.9^-9^) following knockdown, including multiple kinetochore and chromosomal passenger complex components, respectively.

In terms of specific cell cycle regulated genes/proteins, we focused on those that previously registered a significant loss of fitness following knockdown (23) and now with a RIT-seq based cell cycle progression functional assignment. Examples include putative RBPs of the DNA polymerase suppressor 1 (PSP1) family, which display mRNA upregulation in G_1_, protein upregulation in S phase, cell cycle regulated phosphorylation and accumulation in G_2_M following knockdown (see ***Figure 5D*** and ***Figure 7A***). The kinetochore components, KKT1 and KKT7, and also CRK3, all display mRNA upregulation in S phase, protein upregulation in G_2_ and M, cell cycle regulated phosphorylation and accumulation in G_2_M following knockdown (see ***Figure 6D*** and ***Figure 7B***); KKT10 and CYC6 report a similar profile (see ***Figure 6D***), except for the mRNA regulation component. The cytokinesis initiation factors, CIF1 and CIF2, also display mRNA upregulation in S phase, protein upregulation in G_2_ and M and cell cycle regulated phosphorylation, but instead accumulation in the >4C pool following knockdown (see ***Figure 3E***). Finally, the chromosomal passenger complex components, CPC1 and AUK1, as well as furrow localized FRW1, report mRNA and protein upregulation in G_2_M and accumulation in the >4C pool following knockdown (see ***Figure 3E***). Thus, several regulators linked to specific cell cycle progression defects by RIT-seq profiling, are themselves regulated.

## Discussion

Despite intense interest and study (13,15), many cell cycle regulators in trypanosomatids remain to be identified and much remains to be learned about cell cycle control and progression in these parasites. DNA staining followed by flow cytometry is a widely used approach for quantifying cellular DNA content and analysing cell cycle distribution across otherwise asynchronous populations. Here, we combined genome scale loss-of-function genetic screening with flow cytometry in bloodstream form African trypanosomes and identify hundreds of genes required for progression through specific stages of the cell cycle.

Functional annotation of the trypanosomatid genomes will continue to benefit from novel high-throughput functional analyses, and RNAi-mediated knockdown has proven to be a powerful approach for *T. brucei*. RIT-seq profiling provides data for almost every gene and, using this approach, we previously described genome-scale loss-of-fitness data (23). Amongst 3117 knockdowns that scored a significant loss-of-fitness in bloodstream-form cells in that screen (42% of all genes analysed) were genes encoding all 18 intraflagellar transport complex subunits (χ^2^ *p* = 1^-6^), 12 of 13 dynein heavy-chains (χ^2^ *p* = 4^-4^), all 8 TCP-1 chaperone components (χ^2^ *p* = 1^-3^), 27 of 30 nucleoporins (χ^2^ *p* = 2^-7^), all eleven glycolytic enzymes (χ^2^ *p* = 2^-4^) and 30 of 31 proteasome subunits (χ^2^ *p* = 2^-9^). This set also included 18 of 19 kinetochore proteins (χ^2^ *p* = 6^-6^), only later identified as components of this essential complex (17). With this study, we now link many of these genes and many more to specific cell cycle defects following RNAi knockdown. A large number of flagellar protein knockdowns, in particular, yield cells with excess DNA, suggesting that DNA replication typically continues following failure to complete cytokinesis. We identified a number of pathways and protein complexes that impact cell cycle progression, such as glycolysis (G_1_/S transition) and the proteasome (likely G_2_/M transition). We also identify many mRNA binding proteins and protein kinases implicated in control of cell cycle progression. Notably, we link multiple known potential and promising drug targets to cell cycle progression defects, such as glycolytic enzymes (86), the proteasome (87), kinetochore kinases (74,88) and other kinases (89).

Prior cell cycle studies have often focused on trypanosome orthologues of known regulators from other eukaryotes. Since genome-scale profiling is unbiased, it presents the opportunity to uncover divergent as well as novel factors and regulators that impact cell cycle progression. Accordingly, we link many previously uncharacterised and hypothetical proteins of unknown function to specific cell cycle progression defects. Thus, we uncover mechanisms with an ancient origin in a common eukaryotic ancestor and others likely reflecting trypanosomatid-specific biology. We also compared our functional data with cell cycle regulated transcriptome and (phospho)proteome datasets.

The digital dataset provided in ***Supplementary File 1*** facilitates further interrogation and further analysis of the genome-scale cell cycle RIT-seq data. We have also made the data available via an interactive, open access, online data visualization tool, which allows searching and browsing of the data (see ***Figure 2—figure supplement 2***). Comparison with existing and new datasets, including with high-throughput subcellular localisation data (90) www.tryptag.org, should also facilitate future studies. Since high-throughput genetic screens typically yield a proportion of false positive ‘hits’, we do urge some caution, however, in particular where outputs are predominantly generated by a single RIT-seq fragment. On the other hand, there are knockdowns in the current dataset that show specific cell cycle phase enrichment yet fail to register a sufficient enrichment to surpass the thresholds applied above. Considering both of these points, we hope that the digital dataset and the online database will serve as valuable resources. Since other important trypanosomatid parasites, including *Trypanosoma cruzi* and *Leishmania*, share a high degree of conservation and synteny with *T. brucei* (91), the current datasets can also assist and inform studies on other trypanosomatids.

In summary, we report RNAi induced cell cycle defects at a genomic scale and identify the *T. brucei* genes that underlie these defects. The outputs confirm known roles in cell cycle progression and provide functional annotation for many additional genes, including many with no prior functional assignment and many that are trypanosomatid-specific. As such, the data not only improve our understanding of cell cycle progression in these important and divergent pathogens but should also accelerate further discovery. Taken together, our findings further facilitate genome annotation, drug-target prioritisation and provide comprehensive genetic evidence for the protein complexes, pathways and regulatory factors that coordinate progression through the trypanosome cell cycle.

## Materials and Methods

### *T. brucei* RNAi library growth and manipulation

The bloodstream form *T. brucei* RNAi library (22) was thawed in HMI-11 containing 1 μg.ml^-1^ of blasticidin and 0.2 μg.ml^-1^ of phleomycin and incubated at 37°C in 5% CO_2_. After approximately 48 h, six flasks, each containing 2 x10^7^ cells in 150 ml of HMI-11 as above, were prepared; 1 μg.ml^-1^ of tetracycline was added to five of them, while one served as the non-induced control. The cells were grown under these conditions for 24 h and then harvested by centrifugation for 10 min at 1000 g. Cells from each flask were then re-suspended in 25 ml of 1x PBS (pH 7.0) supplemented with 5 mM EDTA and 1 % FBS (“supplemented PBS”), centrifuged again for 10 min at 1000 g, and then re-suspended in 0.5 ml of supplemented PBS. To each cell suspension, 9.5 ml of 1 % formaldehyde in supplemented PBS was added dropwise, with regular vortexing. The cells were incubated for a further 10 min at room temperature and then washed twice in 10 ml of supplemented PBS using centrifugation as above. The cells were finally re-suspended at 2.5 x 10^7^ per ml in supplemented PBS and were subsequently stored at 4°C, in the dark.

### Flow cytometry

Fixed cells, 3 x 10^8^ Tet-induced and 10^7^ uninduced, were centrifuged for 10 min at 1000 g, and re-suspended in 10 ml of supplemented PBS containing 0.01% Triton X-100 (Sigma Aldrich). The cells were incubated for 30 min at room temperature, centrifuged for 10 min at 700 g and washed once in 10 ml of supplemented PBS. The cells were then re-suspended in 4 ml of supplemented PBS containing 10 μg.ml^-1^ of propidium iodide (Sigma Aldrich) and 100 μg.ml^-1^ of RNaseA (Sigma Aldrich), and incubated for 45 min at 37°C, in the dark; cells were subsequently kept on ice and in the dark. Immediately prior to sorting, the Tet-induced cells were filtered (Filcon Cup-type filter, 50 μm mesh, BD™ Medimachine) into 5 ml polystyrene round-bottom tubes (BD Falcon). Cells were sorted using the BD Influx^™^ (Becton Dickinson) cell sorter, with BD FACSort^™^ software, at the Flow Cytometry and Cell Sorting Facility in the School of Life Sciences, University of Dundee. The cells were sorted into pools of <2C (∼8 x 10^5^ cells), 2C (G_1_, 1 x 10^7^ cells), 2-4C (S, 1 x 10^7^ cells), 4C (G_2_M, 1 x 10^7^ cells) and >4C (∼5 x 10^5^ cells) based on their DNA content, and collected into 50 ml Falcon tubes (BD Falcon); sorting time was approx. 4 h. The 2C, 2-4C and 4C sorted samples were then run on a FACS LSR Fortessa flow cytometry analyser for a post-sorting quality check. FlowJo v10 was used for data analysis and visualisation.

### RNA interference target amplification

The five pools of Tet-induced, sorted cells as well as uninduced or induced, but unsorted cells, were lysed overnight at 56°C in 50% (v/v) of Buffer AL (Qiagen) and 0.5 mg.ml^-1^ of Proteinase K (Qiagen), to reverse formaldehyde crosslinking. Genomic DNA was then extracted using the DNeasy Blood and Tissue DNA extraction kit (Qiagen), according to the manufacturer’s instructions, with the exception that each sample was eluted in 50 μl of Buffer AE. The whole sample (range = 140-840 ng) was used for PCR, in a 100 μl reaction, using OneTaq (NEB), and the Lib3F (CCTCGAGGGCCAGTGAG) and Lib3R (ATCAAGCTTGGCCTGTGAG) primers and with the following programme: 94°C for 4 min, followed by 27 cycles of 94°C for 30 sec, 55°C for 30 sec and 68°C for 2 min and 10 sec, and a final extension of 68°C for 5 min. The PCR products were then purified using the Qiaquick PCR extraction kit (Qiagen), as per the manufacturer’s instructions, and eluted in 30 μl of nuclease-free water (Ambion); two columns per sample.

### RIT-seq library preparation and sequencing

Purified PCR products were used for library preparation and sequencing at the Tayside Centre for Genomic Analysis at the University of Dundee. The PCR products were fragmented using a Covaris M220 sonicator (20% duty factor, 75W peak/displayed power, 60 seconds duration – 3 x 20 sec with intermittent spin down step, 18-20°C temperature; resulting in 250-300 bp enriched fragments), and the libraries were prepared using the Truseq Nano DNA Library Prep kit (Illumina). The samples were multiplexed, and sequenced on an Illumina NextSeq 500 platform, on a 150 cycle Output Cartridge v2, paired-end. Each library was run on 4 sequencing lanes. Base call, index deconvolution, trimming and QC were performed in BaseSpace using bcl2fastq2 Conversion Software v2.17.

### RIT-seq data mapping and analysis

The sequencing data analysis pipeline was adapted from (22). The FASTQ files with forward and reverse paired end reads (4 technical replicates for each samples) were concatenated and aligned to the reference genome v46 of *T. brucei* clone TREU927 downloaded from TriTrypDB (28) using Bowtie2 (92), with the ‘very-sensitive-local’ pre-set alignment option. The alignments were converted to BAM format, reference sorted and indexed with SAMtools (93). The quality of alignments was evaluated with Qualimap 2 (94) using the bamqc and rnaseq options. The Qualimap 2 output files were aggregated with MultiQC (95) and inspected. The alignments were deduplicated with the Picard tools package using the MarkDuplicates function (http://broadinstitute.github.io/picard/); to minimise the potential for overrepresentation of the shortest RIT-seq fragments. Alignments with properly paired reads were extracted with SAMtool view using the -f 2 option and parsed with a custom python script to extract the paired reads containing the barcode sequence (GTGAGGCCTCGCGA) in forward or reverse complement orientation. The genome coverage of the aligned reads was extracted from the bam files using bedtools (96) with the -bg option to output bedGraph files. The bedGraph files were visualized with the svist4get python package (97). Read counts for protein coding sequences and associated untranslated regions (where annotated) were determined from the bam files using featureCounts (98) and normalized to Transcripts Per Kilobase Million (TPM). Dimensionality reduction of the G_1_, S and G_2_M TPM values was performed with the radviz algoritm implemented in the pandas python package (99). The bash script containing the analysis pipline, a conda environment specification file for its execution, the python script to extract barcoded reads and a basic usage example are available at GitHub (https://github.com/mtinti/ritseq_cellcycle). Data were subsequently analysed using a GO-slim set and Gene Ontology tools available via tritrypdb.org and visualised using tools available at huygens.science.uva.nl/PlotsOfData.

## Acknowledgements

We thank R. Clark, A. Rennie and M. Lee of the Flow Cytometry and Cell Sorting Facility, which is supported by the Wellcome Trust (097418/Z/11/Z). We also thank L. Glover for advice on RNAi library manipulation, S. Hutchinson for advice on RIT-seq data analysis and J. Faria for fruitful discussions.

## Author contributions

Catarina A. Marques, Conceptualization, Formal analysis, Investigation, Methodology, Writing—review and editing; Michele Tinti, Data curation, visualisation and analysis, Writing—review and editing; Andrew Cassidy, Investigation, Methodology; David Horn, Conceptualization, Formal analysis, Supervision, Funding acquisition, Project administration, Writing—original draft, Writing—review and editing.

## Data and materials availability

High-throughput sequencing data generated for this study have been deposited in the Short Read Archive (SRA) at https://www.ncbi.nlm.nih.gov/sra/PRJNA641153 under primary accession number PRJNA641153.

## Supplementary Figure Legends

**Figure 1—figure supplement 1.**
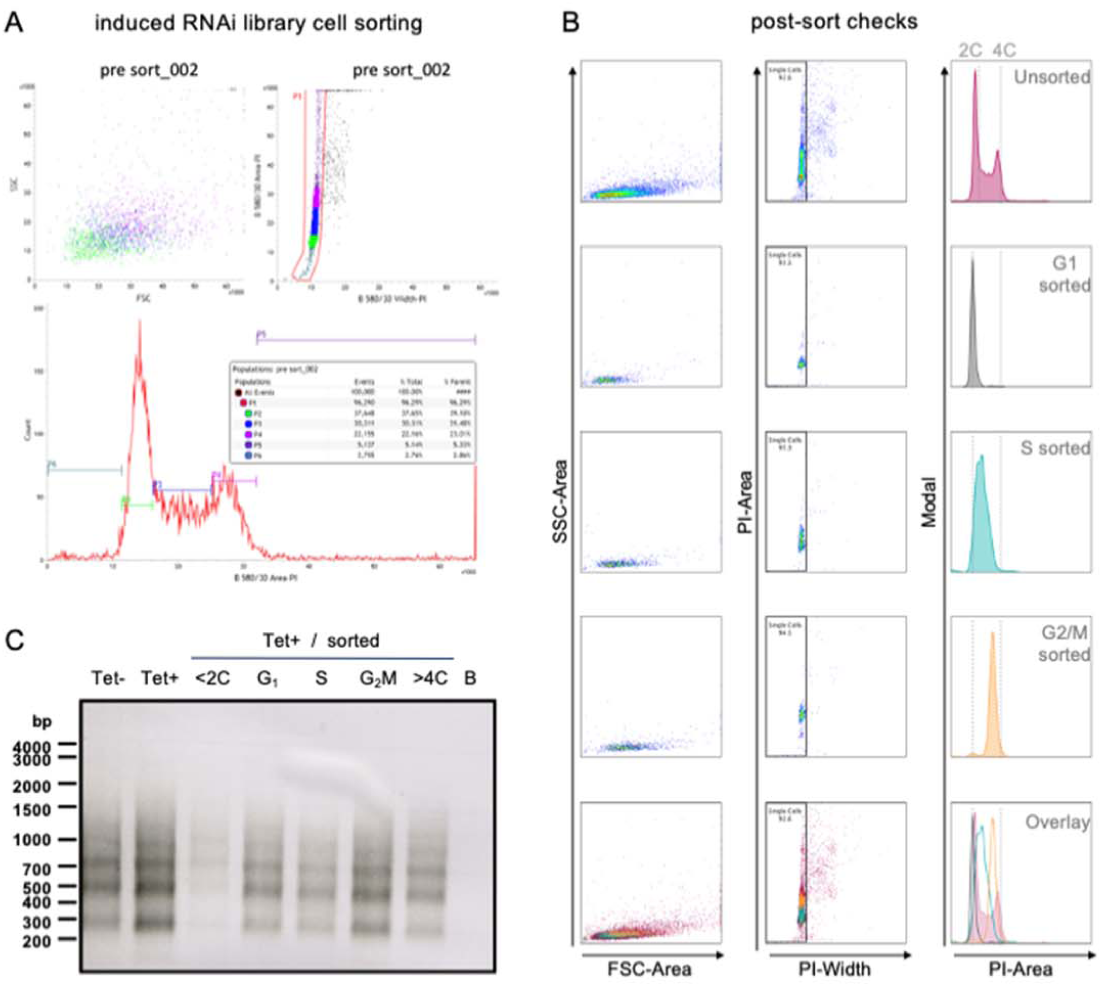
Induced library sorting and RNAi target amplification. (**A**) BD Influx^™^ cell sorter BD FACSort^™^ software workspace data for the sorting session showing the gates and population frequencies. (**B**) Flow cytometry quality control data for the sorted samples. Three types of graphs are shown: SSC-Area x FSC-Area (cell morphology); PI-Area x PI-Width (cells stained with PI; the gate excludes cell aggregates); Modal x PI-Area (cells within the gate set in the PI-Area x PI-Width plot). Top row – unsorted; second row – G_1_; third row – S; fourth row – G_2_M; bottom row – overlay of the sorted and unsorted samples. 2C and 4C refer to unduplicated (diploid genome) and duplicated DNA content, respectively. (**C**) PCR amplification of each sorted sample with Lib3 primers, prior to sequencing.

**Figure 1—figure supplement 2.**
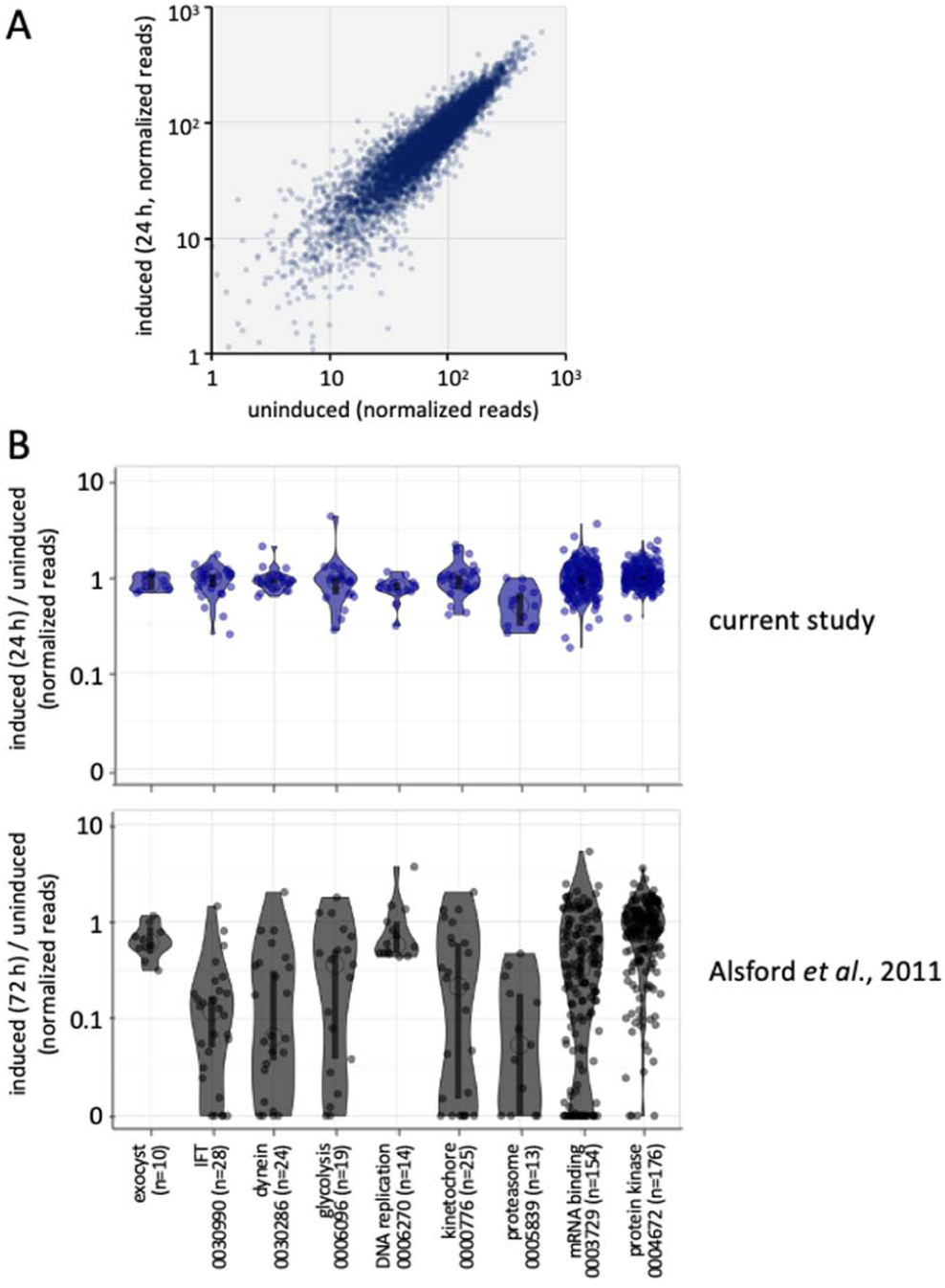
RIT-seq data comparing unsorted libraries. (**A**) The plot shows read-counts for 7,204 genes and for uninduced and 24 h induced samples. Reads for only 0.6% of genes dropped by >3-fold following 24 h of knockdown. (**B**) The violin plots show relative read-counts for cohorts of genes and reflect data distribution. Open circles indicate median values and the vertical bars indicate 95% confidence intervals. Read-counts remain relatively high after 24 h knockdown in the current study, when compared to read-counts after 72 h knockdown in a prior RIT-seq study.

**Figure 2—figure supplement 1.**
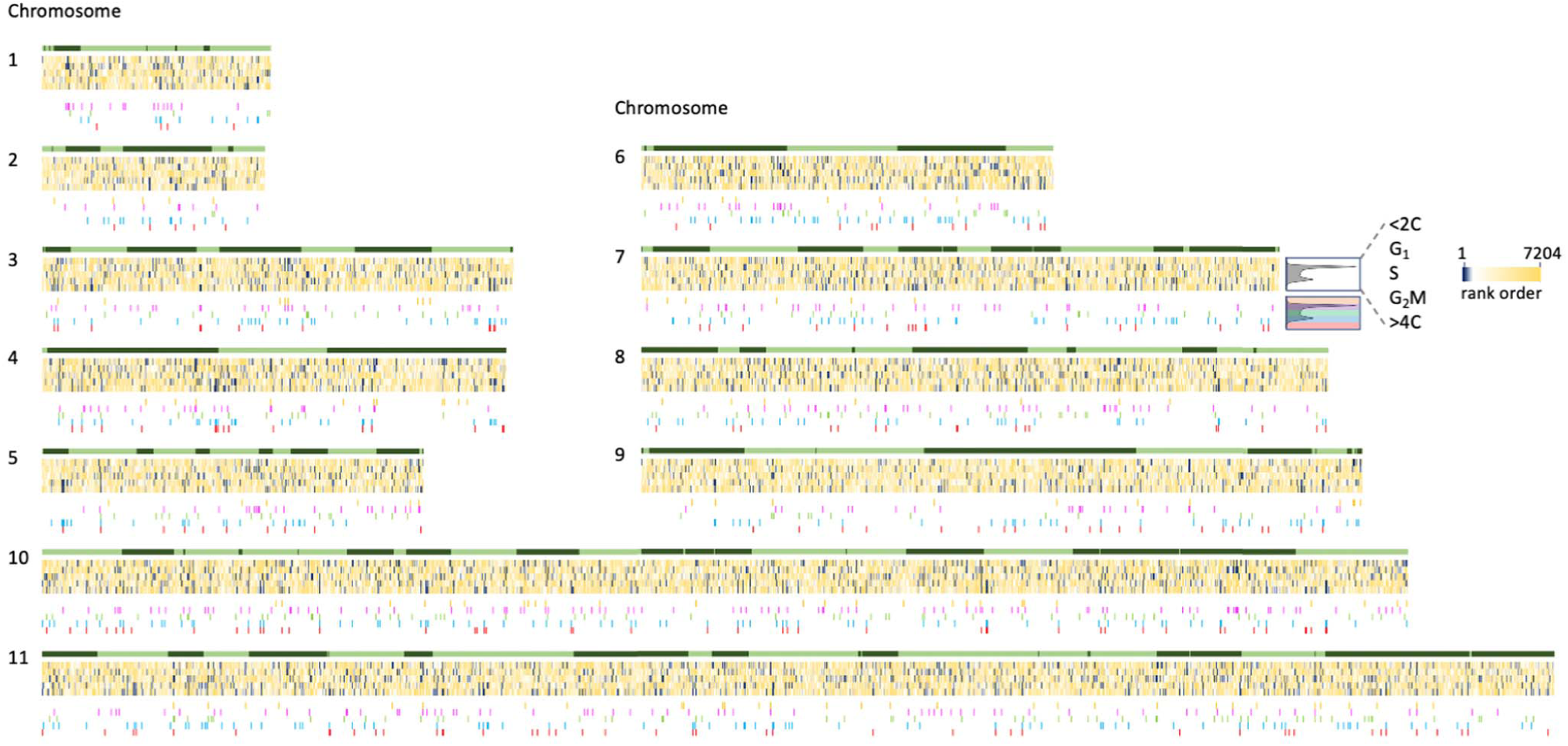
RIT-seq data mapped to *T. brucei* chromosomes; for 7,204 genes and five experiments = 36,020 data-points. Light and dark green indicate polycistronic transcription in the forward and reverse directions, respectively, on each chromosome. The heat maps (dark-blue to yellow) indicate rank enrichment for knockdowns in each cell cycle phase; most enriched are dark-blue. The coloured data-points below indicate those ‘hits’ enriched in each cell cycle phase and used for much of the analysis reported here; 1,158 or 16.1% of genes. See the text for more details.

**Figure 2—figure supplement 2.**
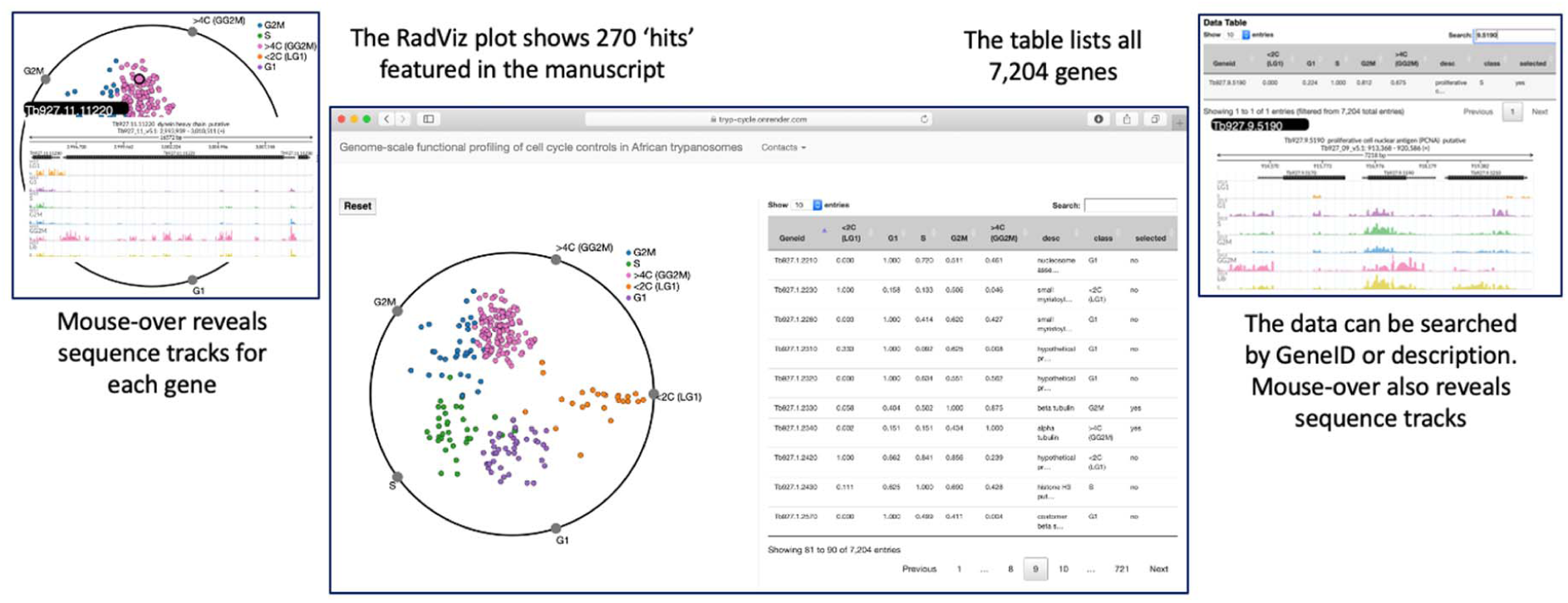
Online RIT-seq data visualization (https://tryp-cycle.onrender.com). In the radial visualization, experimental-points are hours on the clock-face (i.e. related to the angle of the polar coordinate system). The orthogonal axis (i.e. the distance) relates to the relative read-counts across the five experiments. The table on the right shows: Geneid, gene identification number; relative abundance of reads in each sorted sample; desc, gene description; class, the experiment where the gene shows maximum abundance; selected, a binary tag where ‘1’ indicates genes in the radial visualization. Gene coverage images are displayed when hovering over dots on the radial visualization or over table rows. Data transformation to aid visualization: Normalised TPM values from ***Supplementary file 1*** were used, except the <2C and >4C values were divided by the sum of TPM values from all five sorted samples. we elevated the values to the power of 2.1 to maximize differences. We then normalized the values raw wise for each gene, by dividing values by the maximum. The transformed data was then fed to the radial visualization algorithm implemented in D3.js (https://github.com/d3/d3); code and web page were adapted from the repository at https://github.com/WYanChao/RadViz

**Figure 3—figure supplement 1.**
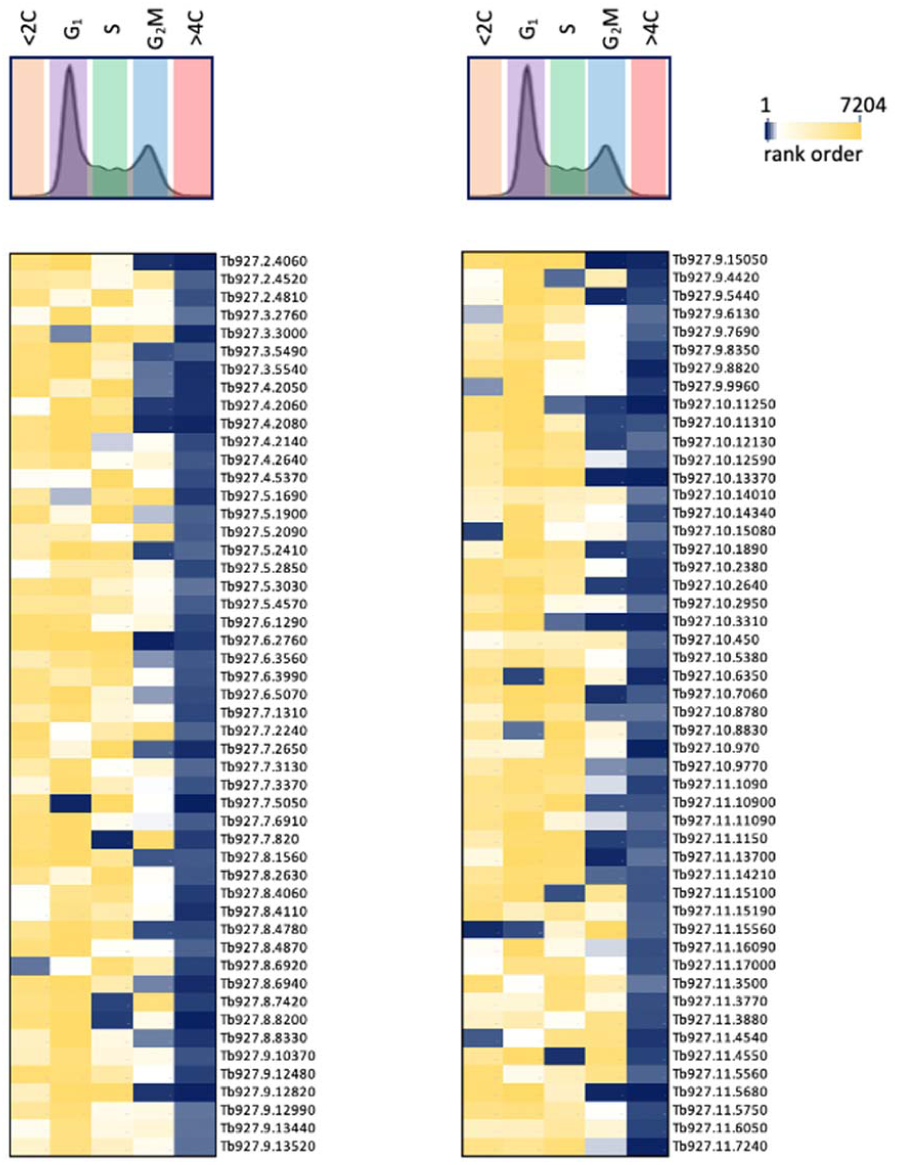
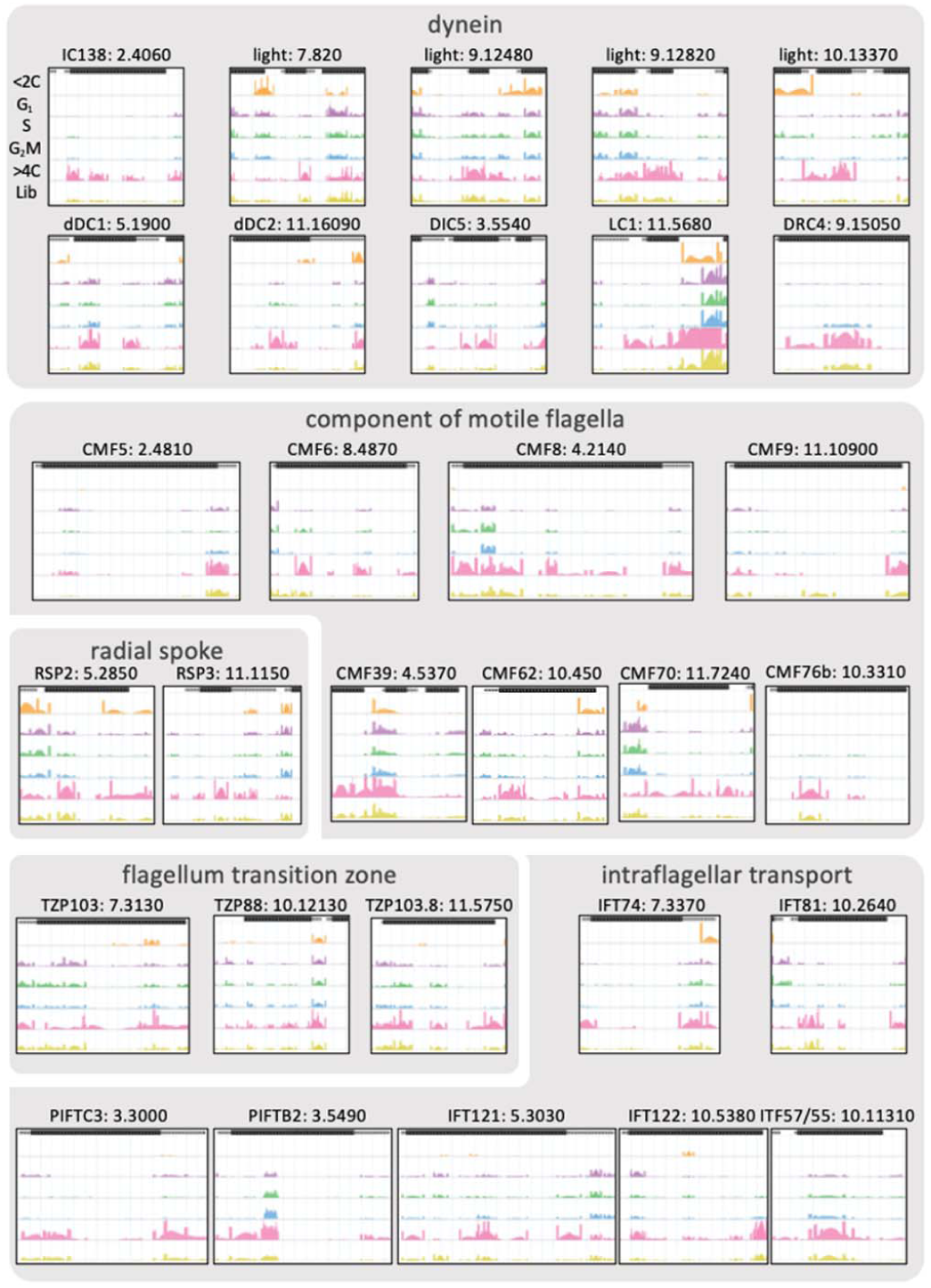
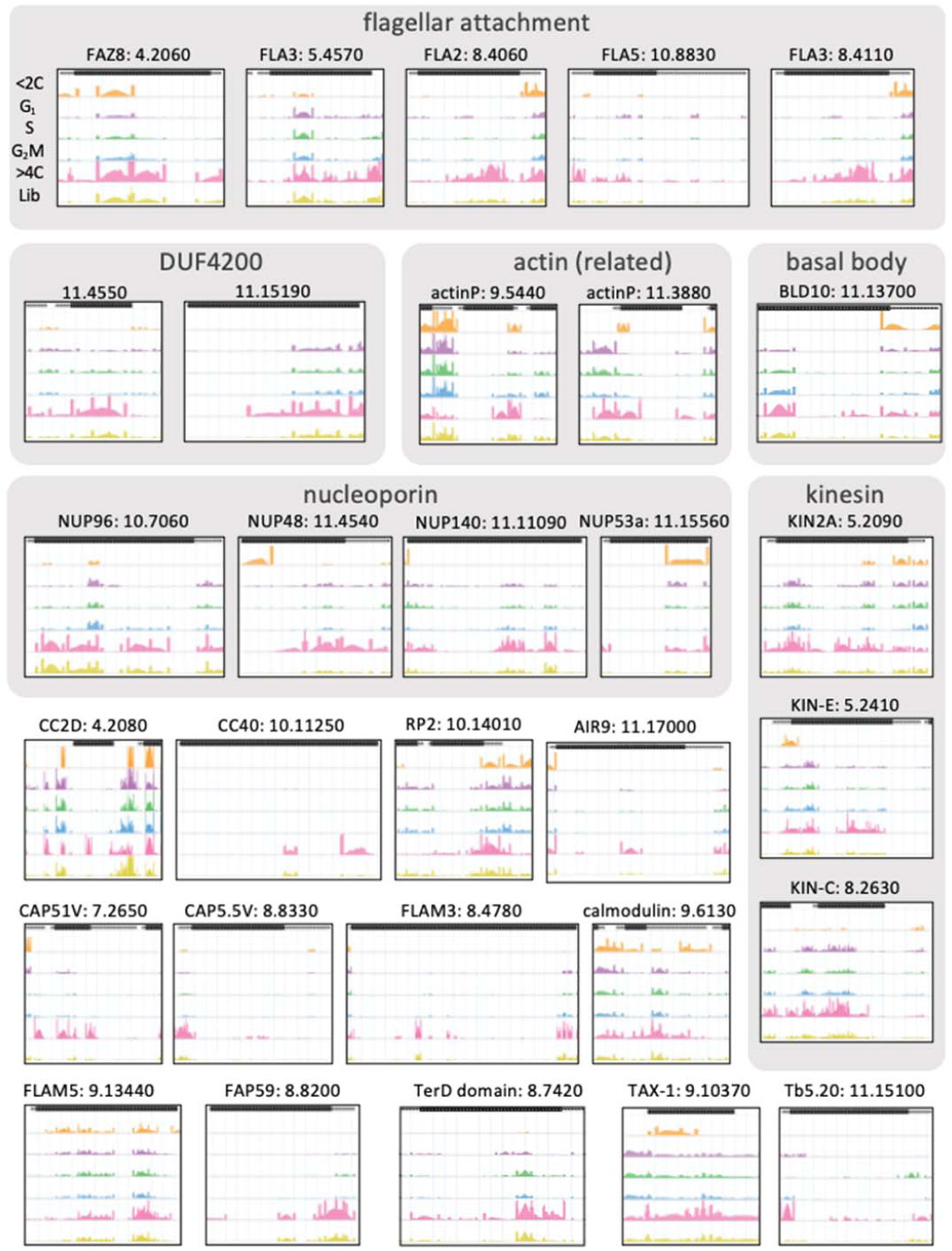
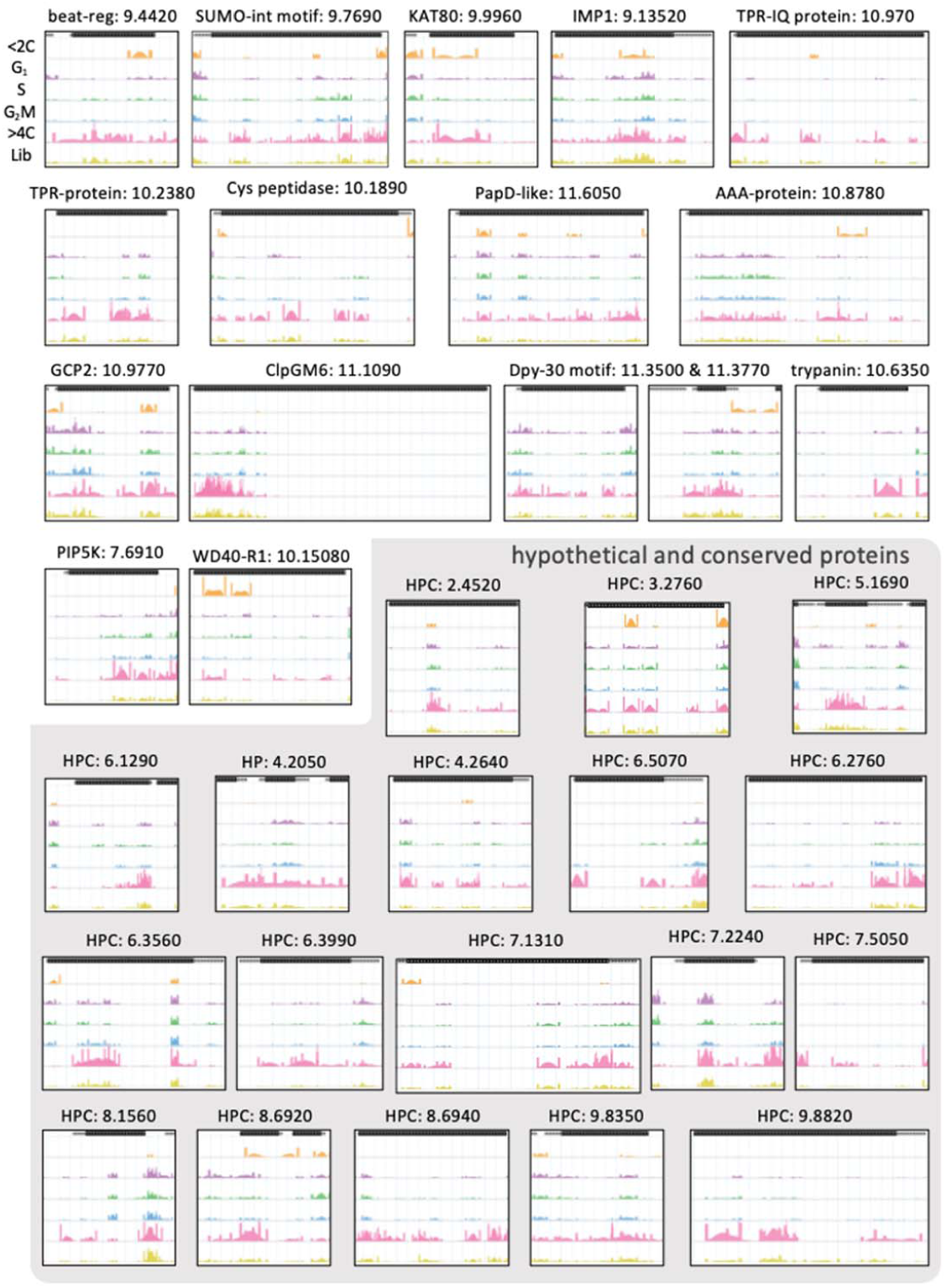
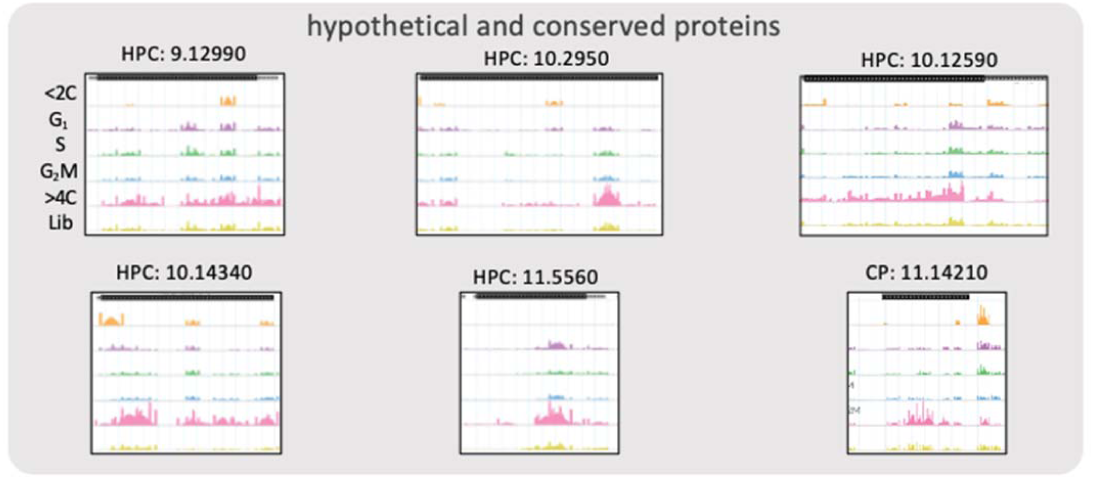
Cytokinesis defects associated with endoreduplication. One hundred example RIT-seq cell cycle profiles are shown for hits overrepresented in the >4C pool. Page 1 shows the heatmaps indicating relative representation in all five sorted pools; blue, most overrepresented. Subsequent pages show read-mapping profiles for each gene; see Figure 1B for further details.

**Figure 7—figure supplement 1.**
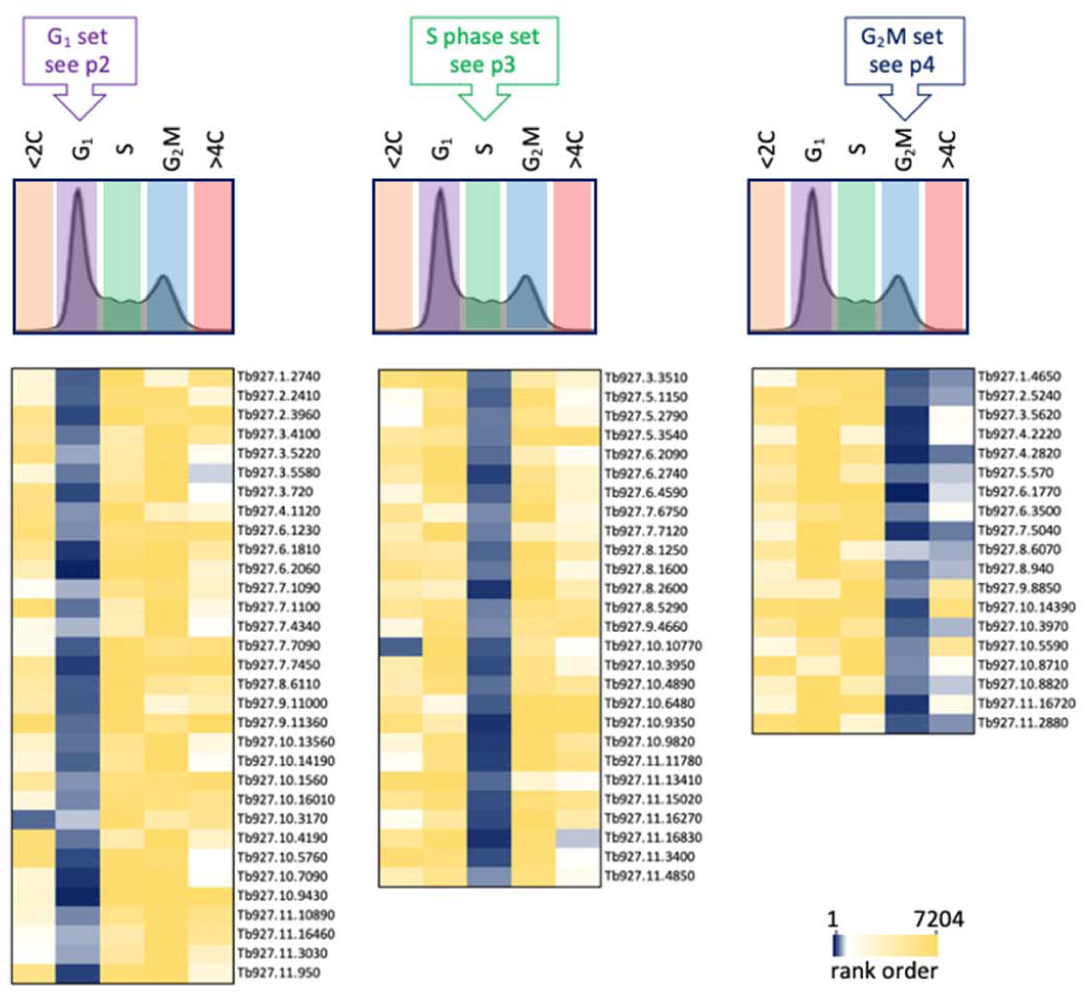
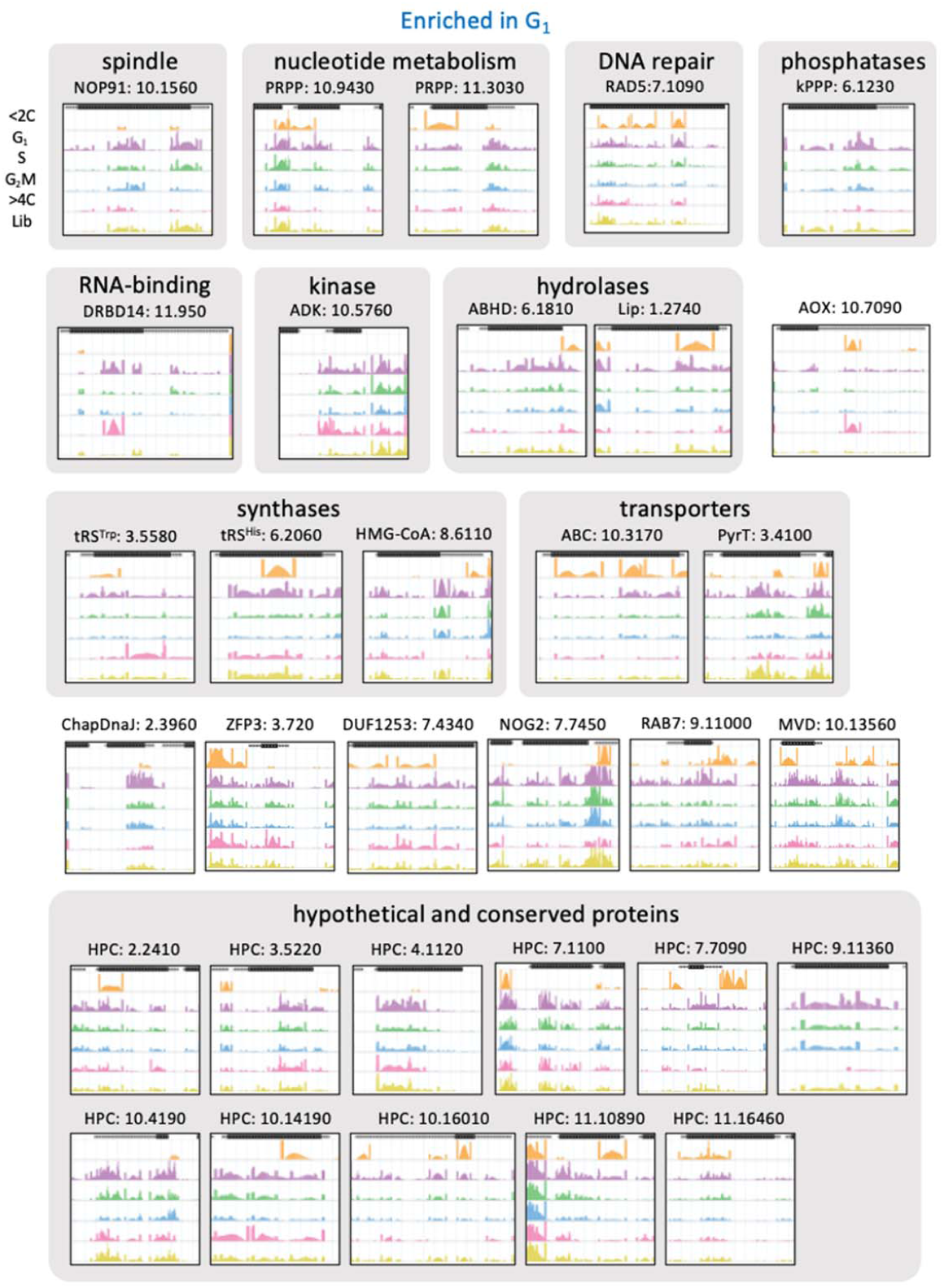
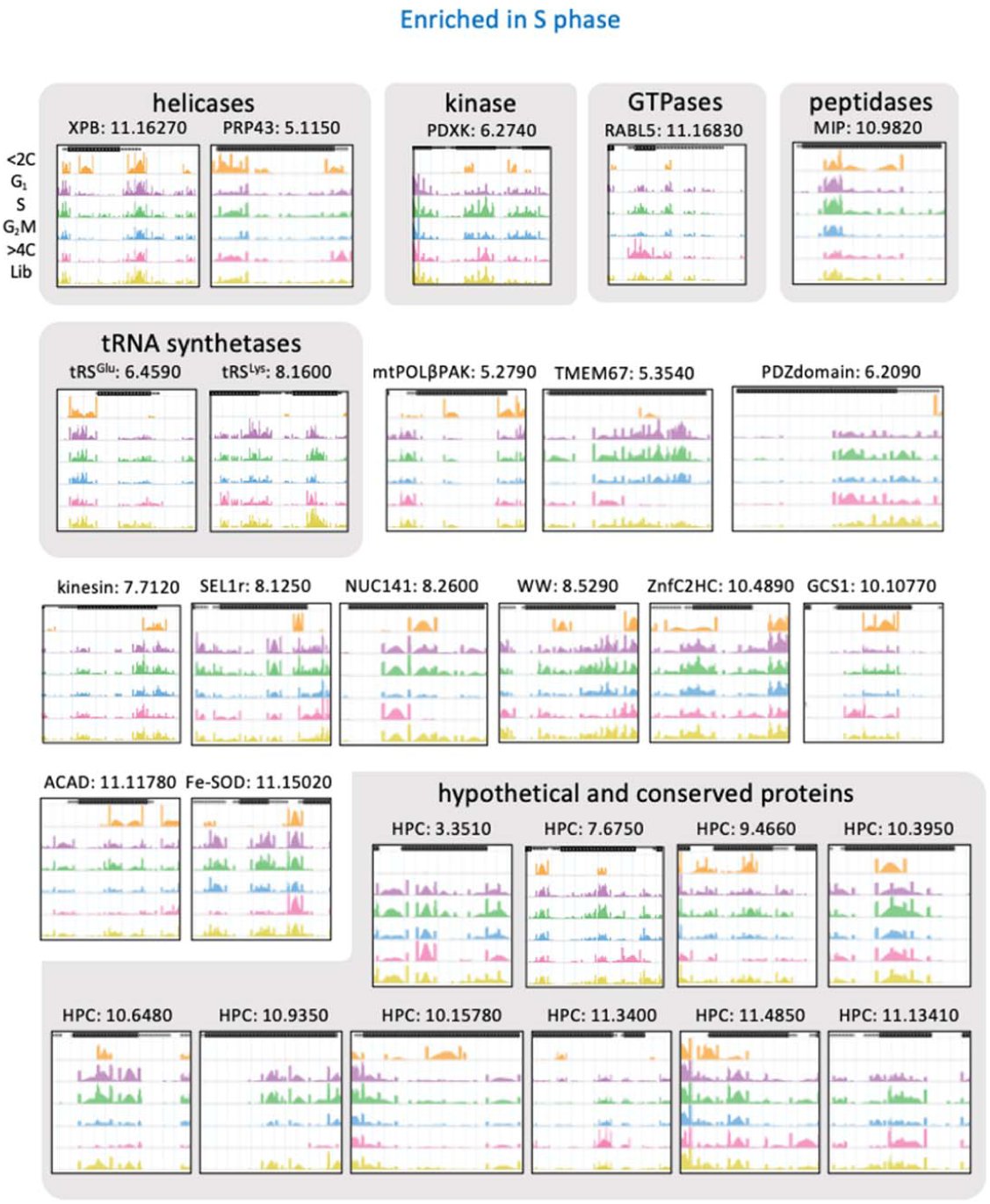
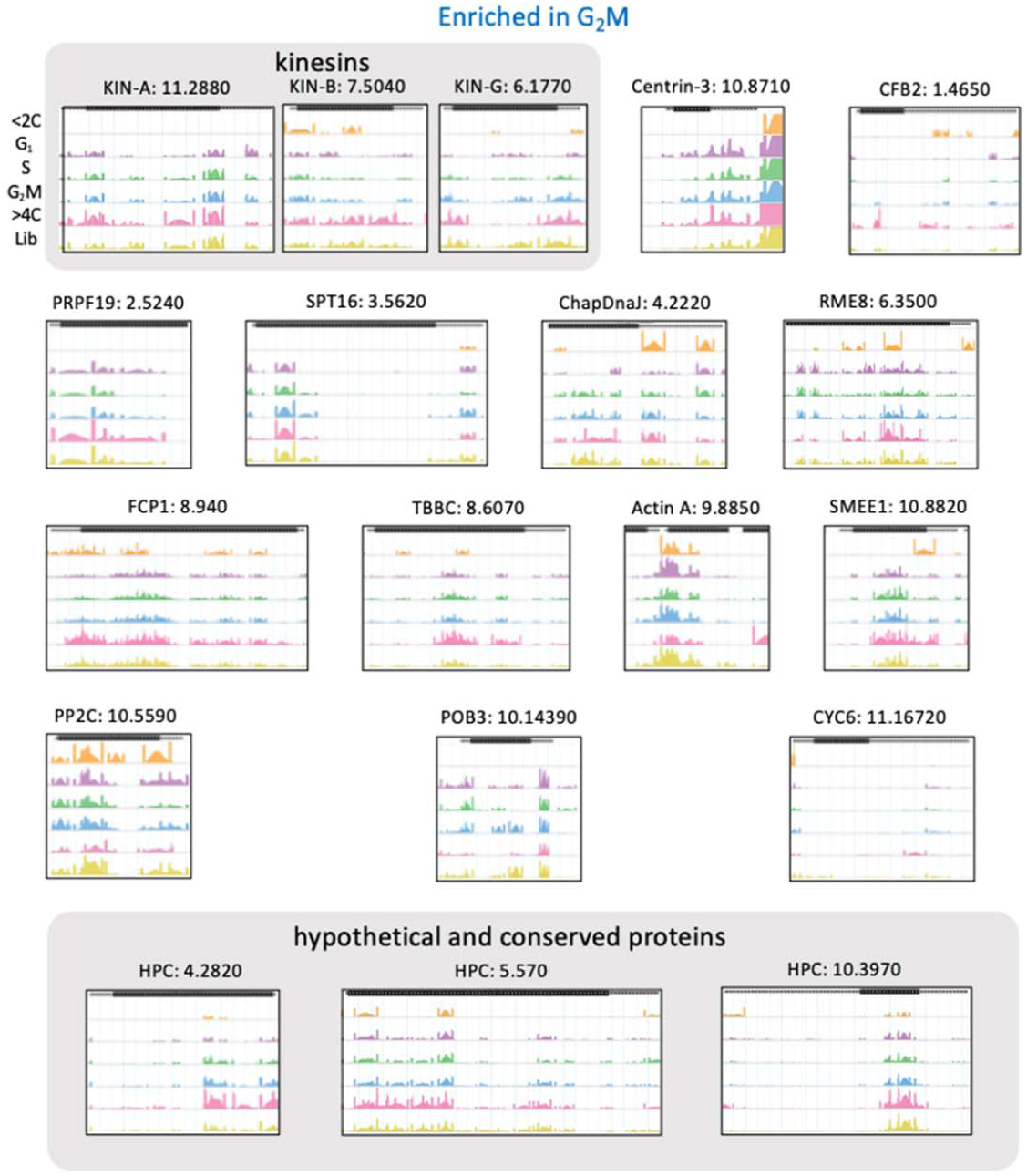
Knockdowns associated with gap and S phase defects. Seventy-eight example RIT-seq cell cycle profiles are shown for hits overrepresented in the G_1_, S phase and G_2_M experiments. Page 1 shows the heatmaps indicating relative representation in all five sorted pools; blue, most overrepresented. Subsequent pages show read-mapping profiles for each gene; see Figure 1B for further details.

***Supplementary file 1.*** RIT-seq digital data. The Excel file reports the total fragment counts for the uninduced (column C) and induced (column D) RNAi libraries, for the five sorted samples (columns E-I), normalised read-counts for the sorted samples (columns J-N) and normalised barcoded read-counts for the G1 (column O), S (column P) and G2M (column Q) samples. The final column (R) indicates Figure numbers for genes shown in the manuscript. TPM, Transcripts Per kilobase Million. Coloured values in column J-Q indicates enrichment in those samples; >1.5-fold the sum of normalised TPM in the G1+S+G2M samples in columns J and N, >41.66% (>25% above the mean) in columns K-M and also >40% (>20% above the mean) in columns O-Q. Genes considered in Figure 2 and used to generate Gene Ontology profiles surpass both the 25% total reads and 20% barcoded reads thresholds i.e. coloured in columns O-Q. Data are presented for 7,204 genes, which is 98% of the non-redundant gene set; all genes register >99 total reads across the five sorted samples.

